# Defects in the HIV immature lattice support essential lattice remodeling within budded virions

**DOI:** 10.1101/2022.11.21.517392

**Authors:** Sikao Guo, Ipsita Saha, Saveez Saffarian, Margaret E Johnson

## Abstract

For HIV virions to become infectious, the immature lattice of Gag polyproteins attached to the virion membrane must be cleaved. Cleavage cannot initiate without the protease formed by the homo-dimerization of domains linked to Gag. However, only 5% of the Gag polyproteins, termed Gag-Pol, carry this protease domain, and they are embedded within the structured lattice. The mechanism of Gag-Pol dimerization is unknown. Here, we use reaction-diffusion simulations of the immature Gag lattice as derived from experimental structures, showing that dynamics of the lattice on the membrane is unavoidable due to the missing 1/3 of the spherical protein coat. These dynamics allow for Gag-Pol molecules carrying the protease domains to detach and reattach at new places within the lattice. Surprisingly, dimerization timescales of minutes or less are achievable for realistic binding energies and rates despite retaining most of the large-scale lattice structure. We derive a formula allowing extrapolation of timescales as a function of interaction free energy and binding rate, thus predicting how additional stabilization of the lattice would impact dimerization times. We further show that during assembly, dimerization of Gag-Pol occurs stochastically and therefore must be actively suppressed to prevent early activation. By direct comparison to recent biochemical measurements within budded virions, we find that only moderately stable hexamer contacts (−12k_B_T<ΔG<-8k_B_T) retain both the dynamics and lattice structures that are consistent with experiment. These dynamics are likely essential for proper maturation, and our models quantify and predict lattice dynamics and protease dimerization timescales that define a key step in understanding formation of infectious viruses.

**Statement of Significance:** For retroviruses such as HIV-1, the Gag polyprotein assembles an immature lattice that ensures successful budding from the cell plasma membrane. The first step in the subsequent maturation requires a pair of protease domains embedded within the lattice to form a homodimer. We show here that this homo-dimerization can proceed within minutes despite involving a small subset of Gag monomers, due to the incompleteness of the immature lattice. Using reaction-diffusion simulations, we quantify timescales of first dimerization events between the protease domains and define a formula to extrapolate across a range of energies and rates. Our models illustrate how protein contacts can be weakened to disrupt lattice assembly or stabilized to slow the remodeling essential for viral infectivity.

## INTRODUCTION

A key step in the lifecycle of retroviruses such as HIV-1 is the formation of new virions that assemble and bud out of the plasma membrane(1, 2). These new virions are initially in an immature state, characterized by a lattice of proteins attached to the inner leaflet of the viral membrane(3, 4). This immature lattice is composed of Gag, Gag-Pol, and genomic RNA (gRNA), with some accessory proteins known to be included for the HIV-1 virion (1). The Gag polyprotein is common to all retroviruses and makes up most of the observed lattice underlying the virion membrane. Within the lattice, 95% of the monomers are Gag (which has 6 domains), and 5% are Gag-Pol, which has the 6-domain Gag followed by protease, reverse transcriptase and integrase domains embedded within the same polyprotein chain(1). The structure of the immature lattice has been partially resolved using sub-tomogram averaging cryotomography(5), revealing Gag monomers that form hexametric rings assembled into a higher-order assembly via additional dimerization contacts. For maturation and infectivity of HIV virions (6, 7), the Gag proteins within the immature lattice must be cleaved by the protease formed from a dimer of Gag-Pol. Importantly, the lattice covers only 1/3 to 2/3 of the available space on the membrane(8, 9). The incompleteness of the lattice results in a periphery of Gag monomers with unfulfilled intermolecular contacts. Recent work showed that these peripheral proteins provide more accessible targets for proteases (10). Here, we address a distinct question on an earlier step in maturation: does the incompleteness of the lattice allow for dynamic rearrangements that ensure that protease domains embedded within the lattice can find one another to dimerize?

Homo-dimerization of the protease domain is necessary for its initial activation (11), with recent cryoEM work demonstrating the dimer can form while attached to Gag-Pols(12). Once activated, the protease triggers a cascade of cleavage reactions, starting with its own(13–15). After the Gag monomers have been cleaved, the newly released domains assemble within the virion cavity to form the HIV mature capsid(11, 16). While the HIV protease has thus been studied extensively(16), the mechanism of its initial activation has not been established. Recent measurements indicate that protease activation can occur within ~100s following assembly of the lattice (17). Here, we use reaction-diffusion simulations of assembled Gag lattices with varying energies and kinetic rates to test how lattice structure and stability can support dimerization of the Gag-Pols at this timescale. We determine which mechanisms of inhibited activation, large-scale lattice remodeling, or dissociation and rebinding of Gag-Pol molecules promote dimerization as an essential step in understanding viral maturation.

Stability of the immature lattice seems to be balanced to promote assembly but also allow for efficient proteolysis and maturation (18); thus mutations and inhibitions that shift the immature lattice stability alter the infectivity of the virus (18). Maturation inhibitors that bind to the immature Gag lattice are thought to stabilize the lattice, preventing cleavage and maturation, which results in loss of viral infectivity (19). Mutations that impede binding of the immature Gag hexamers to inositol hexakisphosphate (IP6) destabilize the lattice, again affecting infectivity (20). The strength of the hexameric contacts is not known, as it is sensitive to co-factors like IP6 and RNA both *in vitro*(21) and *in vivo*(22) (18). The lattice is linked to the membrane via lipid binding and myristolyation (3, 4), and thus the increased concentration on the budded membrane will drive distinct dynamics and stability than those expected in a 3D volume due to dimensional reduction(23, 24). Identifying regimes of binding stabilities and rates that can support assembly and simultaneously support dynamics or remodeling of the immature lattice is thus important for understanding the requirements for forming infectious virions.

With our simulations, we are then prepared to test distinct mechanisms of protease dimerization possible within the immature lattice. Two primary dynamic mechanisms are possible. 1) large-scale remodeling of the lattice could bring together two fragments that contain protease monomers and 2) protease monomers could unbind and reattach at new lattice sites to promote dimerization. Thus, with Gag-Pols incorporated into the lattice at distinct spatial locations, there must therefore be dynamic remodeling of monomers or larger patches within the lattice. Recent experiments show clear evidence of lattice mobility in viruslike particles (VLPs) (25), which are produced by cells expressing only HIV Gag proteins and assemble a Gag lattice very similar to the immature HIV virions(26). Measurements using iPALM microscopy and biochemical cross-linking experiments indicate that Gag lattices that cannot undergo maturation nonetheless exhibit large-scale motion and binding events between individual Gag monomers (25). Furthermore, structural analysis of the immature lattice indicates that the edges of the ~2/3 complete lattice contain Gag monomers that are attached with fewer links(10). These ‘dangling’ proteins would be able to more freely detach and reattach at distinct sites. From such cryoET structures(10), however, it is not possible to measure the dynamics of the lattice and Gag monomers. We note a third mechanism allows the proteases to dimerize during assembly, so they are already adjacent. Experimental evidence indicates they would have to remain as an autoinhibited dimer until after budding occurred, to ensure that the full-length Gag is assembled into budded virions(16). Autoinhibition would prevent early activation of proteases that is known to significantly limit particle formation (27), and cause assembly defects (28). While we cannot test molecular mechanisms of autoinhibition with our model, we can quantify the likelihood of protease dimerization during assembly.

Previous modeling work studying the HIV-1 immature lattice has captured similar structural features to our work but has not interrogated the membrane bound lattice dynamics and their implications for protease dimerization. Coarse-grained molecular scale models of the immature Gag lattice established interaction strengths between Gag domains that are necessary to maintain a hexagonal lattice ordering, as well as changes in structure following mutation(29). Molecular dynamics simulations of incomplete hexamers along the immature lattice gap-edge demonstrated conformational changes in Gag monomers that indicate lower stability and likely targets for protease cleavage(10). Coarse-grained simulations of lattice assembly in solution(30) and on membranes(31) have identified the importance of co-factors, including the membrane, RNA, and IP6 in stabilizing hexamer formation and growth. With our reaction-diffusion approach, we have access to longer timescales despite the large system size (~2500 monomers), and precise control over the association kinetics. We can thus quantify the dynamics and kinetics of the assembled lattice over several seconds for multiple model strengths and rates.

In this work, we initialize Gag monomers into their immature lattices on the membrane, as they would be structured after budding from the host cell but prior to maturation(10). We use reaction-diffusion simulations to both assemble these immature lattices, as well as to characterize the timescales of remodeling and Gag dynamics within the incomplete lattices. We validate that our structured lattices conform to those observed in cryoET, and we verify that the specified free energies and rates of association between our Gag monomers are validated in simpler models. We first characterize the likelihood of the Gag-Pol monomers to dimerize during the assembly process. We find that although they represent only 5% of the monomers that assemble into the lattice, the stochastic assembly will ensure that at least a pair of them are adjacent within the lattice, even if they do not engage in a specific interaction. We next show that, if, on the other hand, the molecules are distant from one another, they would need to detach, diffuse, and reattach stochastically at the site of another Gag-Pol molecule. By modulating the kinetics and energetics of Gag-Gag contacts, we quantify how the overall time for dimerization depends on unbinding, and rebinding, with the 2D diffusion contributing negligibly to the overall time. Lastly, we show how the mobility of the lattice causes binding events that are consistent with biochemical measurements (32), and decorrelation of the lattice that is qualitatively consistent with recent iPALM measurements on immature Gag lattices(25). Our results show that the stochastic dimerization of two Gag-Pol molecules would need to be actively suppressed or inhibited to effectively prevent early activation, and that otherwise, even stable lattices can support Gag-Pol dimerization events due to dynamic remodeling.

## MODELS AND METHODS

### Model components and structural details

Our model contains Gag and Gag-Pol monomers enclosed by a spherical membrane. The membrane contains binding sites for the Gag monomers. The Gag-Pol is structurally identical to the Gag, but represents 5% of the total monomer population to track protease locations within the lattice. The model captures coarse structure of the Gag/Gag-Pol monomers as derived from a recent cryoET structure(33) of the immature lattice (Fig 1A). The key features of our rigid body models are the locations of the four binding sites/domains that mediate protein-protein interactions between a pair of Gag monomers and the Gag-membrane interaction. Each Gag/Gag-Pol contains a membrane binding site, a homo-dimerization site, and two distinct hexamer binding sites that support the front-to-back type of assembly needed to form a ring. When two molecules bind via these specific interaction sites, they adopt a pre-defined orientation relative to one another that ensures the lattice will have the correct contacts, distances between proteins, and curvature (Fig 1). See SI for details on the angle determinations. The Gag monomers bind to the membrane from the inside of the sphere, as would be necessary for budding, and we model this as a single binding interaction that captures stabilization from PI(4,5)P_2_ binding and myristolyation (3, 4). Each reactive site excludes volume from only its reactive partners at a distance *σ*. The dimer site reacts with another dimer site at a binding radius of σ=2.21nm. The MA site binds to the membrane at σ=1nm. The hexamer site 1 binds to hexamer site 2 at σ=0.42nm (Fig 1b). Once reactive sites have bound to one another, they are no longer reactive and no longer exclude volume. Therefore, to maintain excluded volume between monomers throughout the simulation, we introduce an additional dummy reaction between the monomer centers-of-mass (COM). The COM sites exclude volume with a binding radius of σ=2.5nm between all monomer pairs. This is necessary to prevent monomers from unphysically diffusing ‘through’ one another when their reactive sites are fully bound.

**Figure 1.**
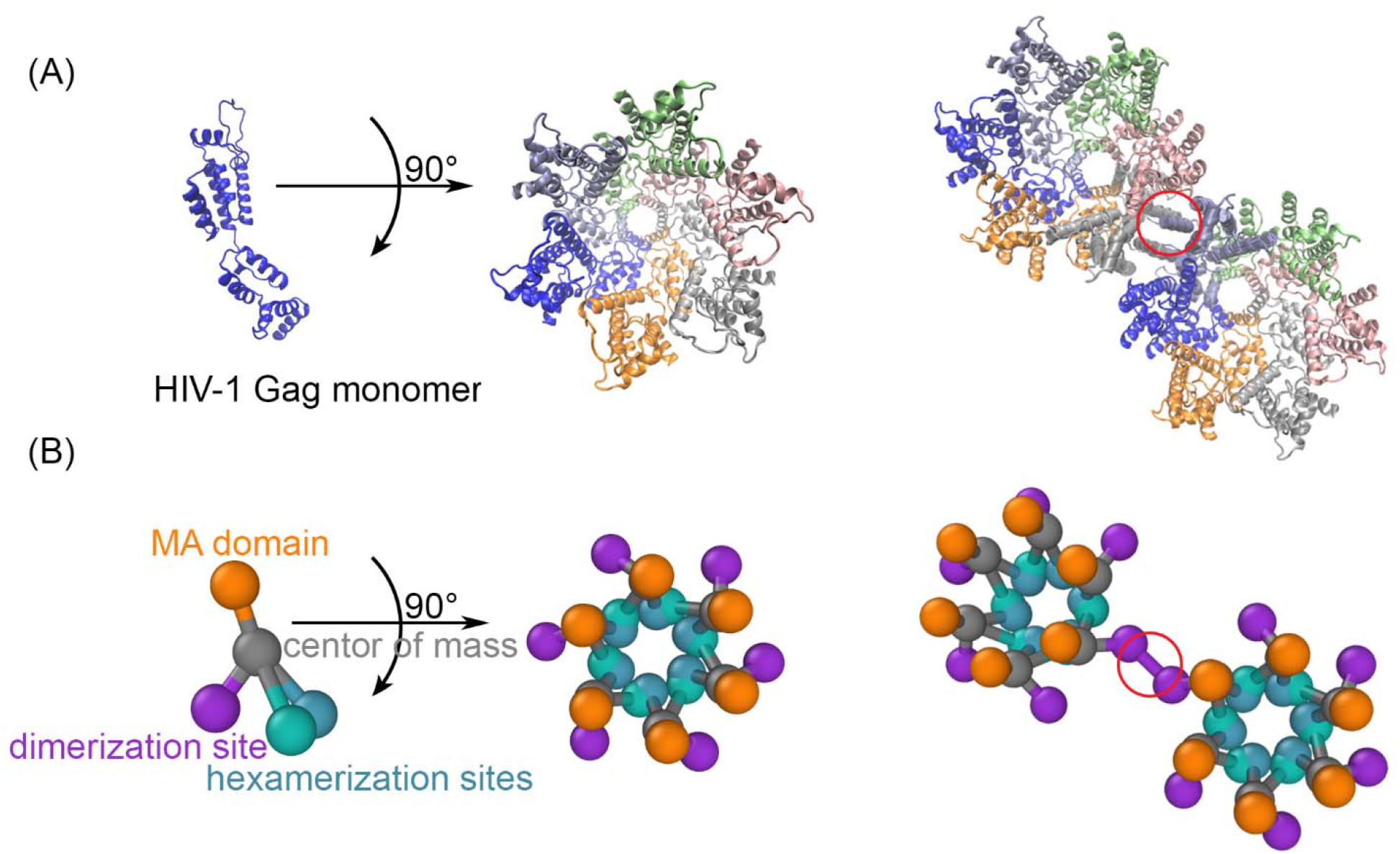
The structure-resolved reaction-diffusion model for Gag assembly on spherical membranes. A) The Gag monomers from the cryoET structure(33) taken from 5L93.pdb is shown on its own, as part of a single hexamer (center), and with a dimerization interface in the red circle that brings together two hexamers (right). B) Our coarse-grained model is derived from this structure to place interfaces on each monomer at the position where they bind. The reaction network contains three types of interactions. The MA domain (orange) binds to the membrane. The position of the MA site is not in the cryoET structure, and we position it to ensure proper orientation of the lattice when bound to the surface. The hexamerization sites (green and blue) mediate the front-to-back binding between monomers to form a cycle. The dimerization site (purple) forms a homo-dimer between two Gag monomers, as illustrated on the right. The reactive sites are point particles that exclude volume only with their reactive partners at the distances shown. Thus the hexamer-hexamer binding radius is 0.42nm, whereas the longer dimer-dimer binding radius is 2.21nm.

### Reaction-diffusion simulations

Simulations are performed using the NERDSS software(34). The software propagates particle-based and structure-resolved reaction-diffusion using the free-propagator reweighting (FPR) algorithm(35). The lipid binding sites on the spherical membrane are modeled using an implicit lipid algorithm, that reproduces the same kinetics and equilibria as the more expensive explicit lipid method(36). We use a time-step Δt=0.2 *μs*. We validated the model kinetics as described in the next section. Software and executable input files for the models used here are provided open source at github.com/mjohn218/NERDSS.

We briefly describe here how the stochastic reaction-diffusion simulations work. Each protein or protein complex moves as a rigid body obeying rotational and translational diffusive dynamics using simple Brownian updates. Each protein binding site is a point particle that can react with a site on another molecule to define the reaction network, as illustrated by the contacts in Fig 1. Reactions can occur upon collisions, with the probability that the reaction occurs evaluated using the Green’s function for a pair of diffusing sites, parameterized by an intrinsic reaction rate *k_a_*, a binding radius *σ*, and the total diffusion of both species(35). This reaction probability is corrected for rigid body rotational motion(37). For proteins that are restricted to the 2D membrane, they perform 2D association reactions with 2D rate constants(38), which are derived from the 3D rate constants by dividing out a lengthscale *h* that effectively captures the fluctuations of the proteins when on the membrane,

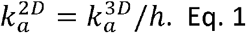

Proteins that do not react during a time-step undergo diffusion as a rigid complex, and excluded volume is maintained for all unbound reactive sites at their binding radius *σ* by rejecting and resampling displacements that result in overlap. All binding events are reversible, with dissociation events parameterized by intrinsic rates *k_b_* that are sampled as Poisson processes. We have that for each reaction, 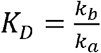, and for the corresponding 2D reaction, we assume the unbinding rates are unchanged, and thus 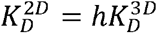.

Binding interactions are dependent on collisions between sites at the binding radius *σ* and are not orientation dependent. Orientations are thus enforced after an association event occurs by ‘snapping’ components into place. Association events are rejected if they generate steric overlap between components of two complexes. Steric overlap is determined using a distance threshold, where here if the distance between molecule centers-of-mass is less than 2.3nm, we reject due to overlap. They are rejected if they generate large displacements due to rotation and translation into the proper orientation, using a scaling of the expected diffusive displacement of 10. Defects ultimately can emerge in the lattice due to the inability of a hexagonal lattice to perfectly tile a spherical surface. These defects result in contacts that are not perfectly aligned; if the contacts are within a short cutoff distance of 1.5σ, they can still form a bond to stabilize the local order, otherwise they are left unbound, weakening the local order.

### Transport

We estimate transport properties from Einstein-Stokes, assuming a higher viscosity for an *in vivo* process. We define D_Gag_(D_Gag-pol_)=10*μ*m^2^/s, D_R,Gag_(D_R,Gag-Pol_)=0.01 rad^2^/s, D_lipid_ = 0.2*μ*m^2^/s. Diffusion slows as complexes grow, consistent with a growing hydrodynamic radius and quantified by the Einstein-Stokes equation. Hence a single protein on the membrane diffuses at 1.96×10^-1^*μ*m^2^/s, and a membrane bound complex containing 1000 proteins diffuses at 1.96×10^-4^ *μ*m^2^/s.

### Energetic and kinetic parameters

We studied lattice dynamics at several strengths defining the free energy Δ*G_hex_* of the hexamerization interaction, at −5.62*k_B_T*, −7.62*k_B_T*, –9.62*k_B_T*, −11.62*k_B_T*, where *k_B_* is Boltzmann’s constant and T is the temperature. We specified the dimerization free energy Δ*G*_dimer_ at −11.62*k_B_T* and −13.62*k_B_T*, which straddles the stability of the measured solution *K_D_* of 5.5 μM or – 12.1*k_B_T* (39). Additional stabilization of the dimer interaction can accompany conformational changes (40), which could drive stronger Gag-Gag binding within the lattice (39). Given the Δ*G* values and using, 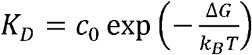, where *c*_0_ is the standard state concentration (1M), we further selected a set of on and off-rates at each free energy, where we used both hexamer and dimer intrinsic binding rates 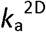 of 2.5×10^-3^ nm^2^/μs, 2.5×10^-2^ nm^2^/μs and 2.7×10^-1^ nm^2^/μs, 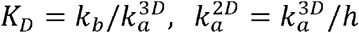. For the Gagmembrane interaction, *h*=2nm, for the Gag-Gag interactions, *h*=10nm, comparable to the size of the Gag monomer.

Our model allows for a strain energy in the formation of closed polygons like hexagons. We set that energy ΔG_strain_ here to +2.3*k_B_T*, meaning that the stability of any closed hexagon within the lattice is slightly lower compared to 6 ideal bonds (i.e. for Δ*G_hex_* =-11.62*k_B_T* it is 5.8 bonds) but still much more stable than a linear arrangement of six Gag monomers which has only 5 bonds. We introduce this penalty to forming closed cycles because of the possibility of imperfect alignment of molecules in the lattice.

We validated the kinetics and equilibrium of our model as it assembled on the membrane when we set the hexamer rates to zero, so it formed purely dimers (Fig S1), and when we set the dimer rates to zero, so it formed purely hexamers (Fig S2). The observed kinetics and equilibria were compared to solutions solved using the corresponding system of non-spatial rate equations, showing very good agreement with apparent intrinsic rates that systematically accounted for excluded volume and the criteria used for accepting association events (Fig S1-S2). Thus, all of the rates and free energies reported in the paper agree with the kinetics and equilibria observed and expected for the sets of binding interactions that make up the full lattice system.

### Simulations for constructing the initial lattices on the membrane

The HIV lattice is composed of Gag and GagPol bound to the inner leaflet of the lipid membrane. To study the remodeling dynamics, we must construct the initial configurations where the lattice is assembled such that it has a specific coverage of the surface (67% or 33%), and is linked to the membrane via lipid binding. We define the membrane sphere of radius 67nm to represent the membrane surface(41). PI(4,5)P_2_ is populated on the membrane surface at a concentration 0.07nm^-2^, and is treated using an implicit lipid method. Thus while binding events reduce the free population of PI(4,5)P_2_, we do not have to propagate the thousands of individual sites. Instead of dumping all Gag monomers in as a bulk unit, we titrate in the Gag and Gag-Pol monomers at a rate of 6×10^-5^ M/s and 3×10^-6^ M/s respectively, which can ensure a ratio of Gag: Gag-Pol of ~ 20:1, consistent with experiment(41). Gag(Gag-Pol) molecules can bind in solution (3D), to the membrane (3D to 2D), and when on the membrane (2D). We forbid the binding between Gag-Pol – Gag-Pol pairs to suppress the ‘activation’ events that could therefore occur during assembly by setting those rates to zero during these simulations. Assembling the Gag monomers into a single spherical lattice is non-trivial due to the size of the lattice and its propensity to nucleate multiple lattice fragments. While the titration of the monomers reduced multiple nucleation events for the membrane system, we found that the easiest way to form a single lattice was by assembling the structure fully in solution, in a volume of (250nm)^3^. The Gag model was otherwise the same, except the rates of dimerization and hexamerization were both set to 10nm^3^/μs, and binding was irreversible. We then put the assembled single lattice into a spherical system by linking this structure to the membrane using one PI(4,5)P_2_ attachment per monomer. We generated 16 initial configurations for each coverage area (67% and 33%). Some initial configurations are shown in Fig. 2.

**Figure 2.**
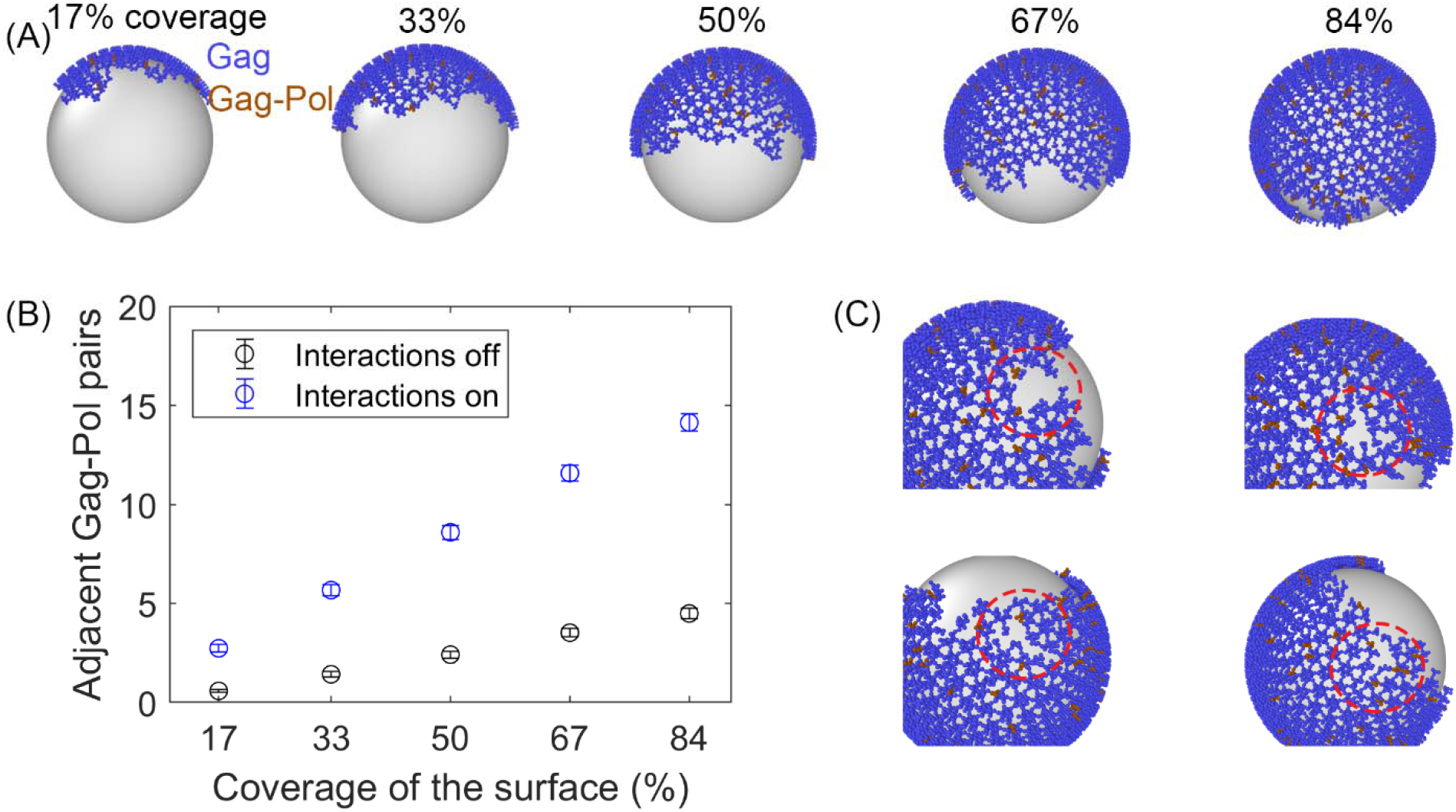
Initial Gag immature lattices within the membrane are assembled via simulation. A) The starting Gag immature lattices are assembled from NERDSS simulations with irreversible binding. ~5% of the monomers are Gag-Pols shown in brown. We note that the silver spheres are shown here only to improve visualization of just one side of the lattice; Gag proteins are attached to the *inner* surface of the budded spherical membrane, consistent with experiment. B) The number of adjacent pairs of Gag-Pol in the initial immature lattice increases with more surface coverage. Even with explicit Gag-Pol to Gag-Pol interactions turned off (black circles), they still end up adjacent to one another. C) Formation of the lattice produces defects that are like those present in cryoET structures.

### Simulations for lattice remodeling dynamics

For each initial configuration we have generated, we perform 6 independent trajectories. See Movie S1 for one trajectory. We perform these 96 simulations for each set of model parameters to generate statistics both within and across initial configurations. For some simulations, fragments of the lattice become sterically overlapped with one another, due to the high density and the time-step size. While this could be eliminated by lowering the time-step, we instead keep the more efficient time-step, and discard these simulation traces which produce overlap. We finally analyze 60 remodeling traces for each parameter set. All the simulation parameters are listed in Table 1. The number of monomers is fixed for each simulation by the initial configuration, so that only binding, unbinding, and diffusion can occur throughout the simulation. During lattice construction, in one set of simulation we set all Gag-Pol to Gag-Pol binding interactions to zero (Fig 2B). Now for the remodeling dynamics, we allow all interactions involving Gag-Pol, and at rates that are identical to those involving Gag, meaning there is no difference between the types except for in their label. The Gag(Gag-Pol) molecules are allowed to diffuse on the membrane, where they can unbind from a molecule and rebind to another with the specified binding rates. Each monomer can also unbind and rebind to the membrane. However, dissociation to solution is extremely rare, as it requires that all Gag monomers in an assembled complex unbind from their lipid before any of the sites rebind.

**Table 1.**
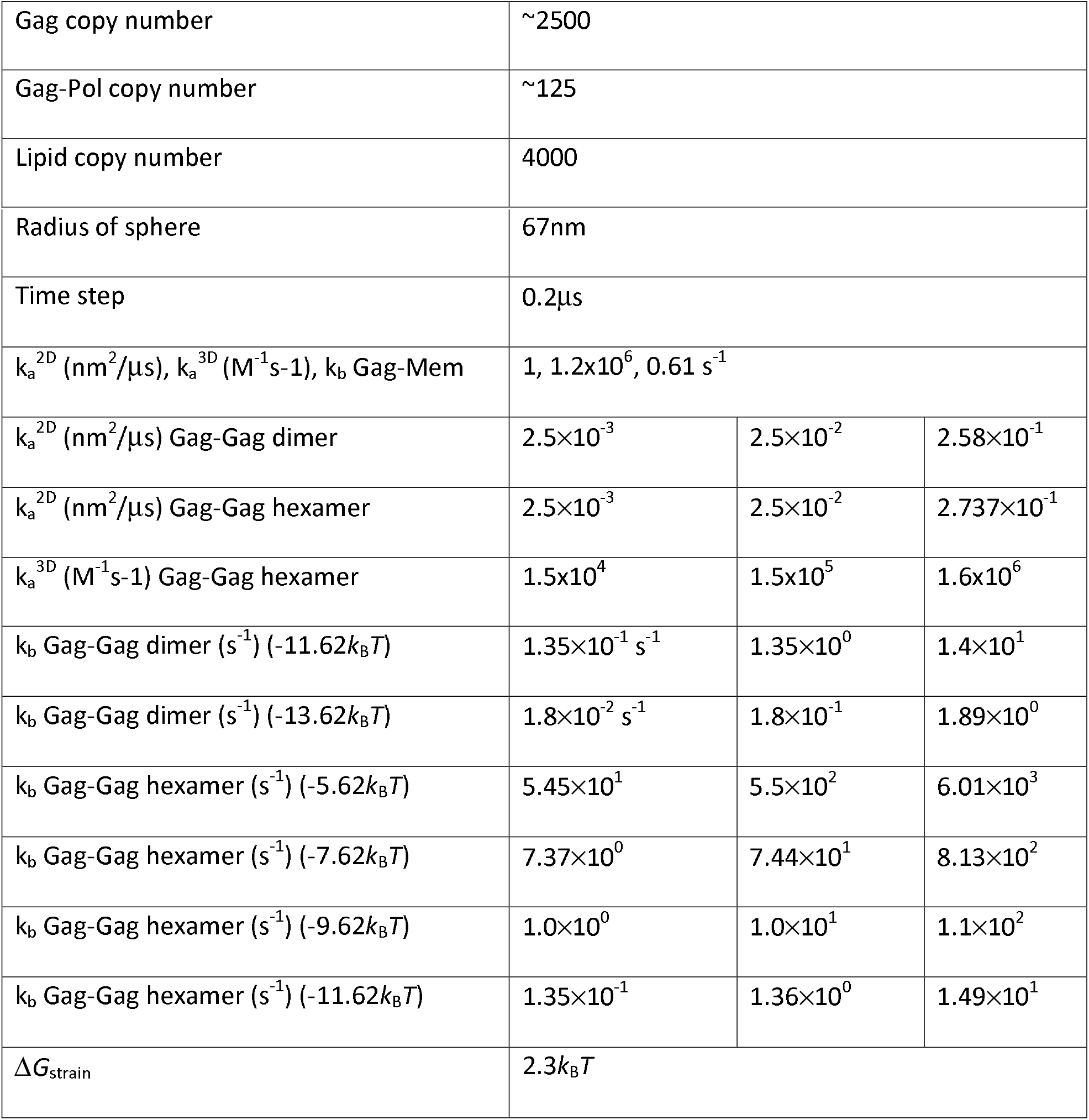
Simulation parameters.

|  |  |
| --- | --- |
| Gag copy number | ~2500 |
| Gag-Pol copy number | ~125 |
| Lipid copy number | 4000 |

|  |  |  |  |
| --- | --- | --- | --- |
| Radius of sphere | 67nm |  |  |
| Time step | 0.2μs |  |  |
| $k_a^{2D}$ (nm<sup>2</sup>/μs), $k_a^{3D}$ (M<sup>-1</sup>s<sup>-1</sup>), $k_b$ Gag-Mem | 1, 1.2x10<sup>6</sup>, 0.61 s<sup>-1</sup> | | |
| $k_a^{2D}$ (nm<sup>2</sup>/μs) Gag-Gag dimer | 2.5x10<sup>-3</sup> | 2.5x10<sup>-2</sup> | 2.58x10<sup>-1</sup> |
| $k_a^{2D}$ (nm<sup>2</sup>/μs) Gag-Gag hexamer | 2.5x10<sup>-3</sup> | 2.5x10<sup>-2</sup> | 2.737x10<sup>-1</sup> |
| $k_a^{3D}$ (M<sup>-1</sup>s<sup>-1</sup>) Gag-Gag hexamer | 1.5x10<sup>4</sup> | 1.5x10<sup>5</sup> | 1.6x10<sup>6</sup> |
| $k_b$ Gag-Gag dimer (s<sup>-1</sup>) (-11.62k<sub>B</sub>T) | 1.35x10<sup>-1</sup> s<sup>-1</sup> | 1.35x10<sup>0</sup> | 1.4x10<sup>1</sup> |
| $k_b$ Gag-Gag dimer (s<sup>-1</sup>) (-13.62k<sub>B</sub>T) | 1.8x10<sup>-2</sup> s<sup>-1</sup> | 1.8x10<sup>-1</sup> | 1.89x10<sup>0</sup> |
| $k_b$ Gag-Gag hexamer (s<sup>-1</sup>) (-5.62k<sub>B</sub>T) | 5.45x10<sup>1</sup> | 5.5x10<sup>2</sup> | 6.01x10<sup>3</sup> |
| $k_b$ Gag-Gag hexamer (s<sup>-1</sup>) (-7.62k<sub>B</sub>T) | 7.37x10<sup>0</sup> | 7.44x10<sup>1</sup> | 8.13x10<sup>2</sup> |
| $k_b$ Gag-Gag hexamer (s<sup>-1</sup>) (-9.62k<sub>B</sub>T) | 1.0x10<sup>0</sup> | 1.0x10<sup>1</sup> | 1.1x10<sup>2</sup> |
| $k_b$ Gag-Gag hexamer (s<sup>-1</sup>) (-11.62k<sub>B</sub>T) | 1.35x10<sup>-1</sup> | 1.36x10<sup>0</sup> | 1.49x10<sup>1</sup> |
| $\Delta G_{\text{strain}}$ | 2.3k<sub>B</sub>T | | |

### Calculation of First-passage times (FPT)

Our primary observable is how long it will take for the first dimerization of Gag-Pols. The ‘clock’ is started from the initialized lattices shown in Fig 2. We note that two Gag-Pols have a chance to be adjacent at the initial configuration (Fig 2), but we ignore these events since dimerization of Gag-Pol before viral release has been experimentally shown to result in loss of Pol components from the virions (42).

### Fitting of the Mean First Passage Times

To derive a phenomenological expression that captures our measure mean first-passage times (MFPT), we used a single global formula for all our model results:

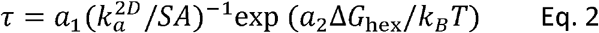

Where there are two fit parameters, *a*_1_ and *a*_2_ and the surface area *SA* – *4πR*^2^ of the sphere is the same for all models. This expression is inspired by characteristic timescales for bimolecular association, which are inversely dependent on the reaction rate (here 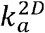)(43). The *SA* is present to ensure the correct units for *τ*, and we empirically observe the relationship between the hexamer free energy and the measured MFPT. We optimized the parameters *a*_1_ and *a*_2_ busing nonlinear fitting in MATLAB to the ln(*τ*) functional form of Eq. 2.

### Calculation of biochemical SNAP-HALO binding timescales

Recent measurements on Gag VLPs quantified dimerization between sub-populations of Gag molecules within the lattice(32). Dimerization events were identifiable because one sub-population of Gag monomers carried a HALO tag, and another carried a SNAP tag, and the addition of a linker HAXS8 produced a covalent linkage of HALO-HAXS8-SNAP. The concentration of these HALO-HAXS8-SNAP structures was then quantified vs time. We therefore reproduced this experiment via analysis of our simulation trajectories. We defined a population of our Gag monomers randomly selected to have a ‘SNAP tag’, and a population randomly selected to have a ‘HALO tag’. For each trajectory, the tagged populations of each were either 5,10, 20 or 40% of the total monomers, to match the experimental measurements (32). We then monitored the number of dimerization events that occurred as a function of time. A dimerization event required that a monomer with a SNAP-tag and a monomer with a HALO-tag encountered one another at a distance less than 3nm (44), where one and only one of these partners must have the covalent linker attached. We randomly selected half of the population of SNAP-Gags to have a linker attached, and half of the population of HALO-Gags to have a linker attached, and thus some encounters between a SNAP and HALO Gag were not productive if zero or 2 linkers were present. Ultimately, however, all dimers could be formed given the symmetric populations containing linkers. In the SI, we provide additional justification for these assumptions based on the permeability of the linker and its biochemical binding strengths (44). Since our simulations are ~20s, we did a linear fit of the last second of the dimer forming kinetics to extrapolate the dimer copies at 3 minutes, which can be used for comparison with the experiment. This extrapolation therefore assumes that dimer formation does not slow down, which it almost certainly does. Therefore, our observables provide an upper bound on the expected number of HALO-SNAP dimers.

### Calculation of number autocorrelation function (ACF)

We calculated an autocorrelation function (ACF) of lattice dynamics to track collective motion comparable to iPALM experiments on Gag virus-like particles (VLPs). Specifically, we counted the number of Gag monomers found on each fixed 1/8 of the sphere surface. The copy numbers within each of the 8 quadrants vary due to diffusion of the lattice, while the total copy numbers on the surface is fixed. The copy numbers at a time point *t* are denoted by *G*(*t*), and thus the autocorrelation function for each quadrant is given by:

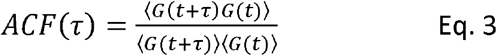

These ACF measurements are comparable to fluorescence correlation spectroscopy measurements (45). For comparison, the ACF for *G*(*t*) across the full sphere surface is 1 at all times, because there is no change in total copy numbers. At long times, when the counts are uncorrelated, this function will go to 1, because 〈*G*(*t* + *τ*)*G*(*t*)〉 → 〈*G*(*t* + *τ*)〈〉*G*(*t*)〉. Furthermore, if the copies are all well-mixed across the surface, then we expect minimal deviations from 1 across all times given these relatively large viewing regions (1/8 of the surface), because there is no source of correlation between copies. As τ→0, the deviation of this ACF from 1 reports on the variance of the fluctuations in the numbers per patch relative to the mean, or the coefficient of variation (CV) squared: 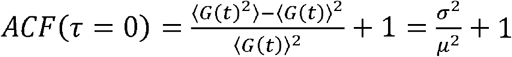. We calculated the number correlation for each quadrant of one simulation trace. We then took the average of all 60 traces for each parameter set. We also used an ensemble averaging approach, where we calculated the copy number correlations and means across multiple quadrants and trajectories before averaging to get the numerator and denominator of Eq. 3. This method provides more statistics on longer time-delays, and is based on assuming that all trajectories are sampling from the same equilibrium distribution. The trends are the same as those we report but shifted up to slightly higher amplitudes before decaying to 1. We note that our ACFs are not truly reporting on equilibrium fluctuations, as we show below that there is clearly some time-dependent changes to the lattice structure that is not reversible due to fragmenting along the edge compared to the initial structures. However, we compared the ACFs calculated for the first half of our simulations vs the last half, for example, and all of the same trends are preserved.

In addition to directly calculating this number autocorrelation, we also sampled it using a stochastic localization approach that directly mimics the experimental measurement (25). For this approach, for each time-point we ‘activate’ a single Gag monomer across the full lattice with a probability p_act_. So for each frame, either 1 or 0 Gag monomers is visible. We identify the quadrant for that monomer, and thus each quadrant produces a sequence of 1s and 0s. After a monomer is activated, it is then bleached, and cannot be localized again. The sequence of localizations is then used to calculate the same ACF, where we use a binning method(45), as in the experimental analysis, to improve statistics on the signal at larger time delays. The agreement between the stochastic measurement of the ACF and the direct measurement of the copy numbers ACF are excellent (see below). We use this stochastic localization method so that we can introduce additional sources of correlation to our measurement of the simulated lattice dynamics (SI Methods), since these measurement artifacts can appear in the real experimental system.

### Analysis of autocorrelation function (ACF) from experimental data on VLPs

The iPALM experiments to characterize lattice dynamics in virus-like particles (VLPs) were previously described and published (25). We describe the analysis of the stochastic localization experiments here because we focus on analyzing a shorter part of the measurement. We analyzed only the first 5000 frames of the measurement, or 500 seconds, because after that time the laser intensity was changed. Based on data collected on 25 VLPs, we analyzed only the VLPs where they reported a large enough fraction of localization events to indicate a reliable measurement, so we included only VLPs where >75% of the quadrants had more than 250 localizations, leaving 11 VLPs. We used the same algorithm (45) as applied to the simulation data to quantify the ACF from the time-dependent sequences of localization events (typically a series of 1s and 0s, with occasionally 2 events per frame). We also calculated the background signal for each VLP. That is, we merged the localization traces from the 8 quadrants into the single localization trace across the entire VLP surface, as the full surface was visualized once per experiment, with localization events then assigned to quadrants. This background signal reports on fluctuations in the total copy numbers of Gag on the surface, which we would expect to be 1 given a perfect measurement, but which was always higher than this due to measurement noise. We divided out this background signal from the averaged signal across the 8 quadrants per VLP, as the ACF would then more accurately report on only the relative change in copy numbers per quadrant (see SI for more details). We then averaged these signals across the VLPs. We performed the same analysis on the 25 VLPs that had been modified by a fixative, first filtering out the VLPs that had too few localization measurements (leaving 13 VLPs), and then otherwise proceeding with the analysis in an identical fashion.

## RESULTS

### A. Assembled lattices on the membrane are structurally similar to those present in cryoET

We were able to assemble a variety of spherical Gag lattices that grew from monomers to a single sphere with our targeted coverage of the surface (Fig 2). The parameters needed to assemble these structures (Methods) were not expected to be representative of the physiologic values, given that they were not assembled as a growing virion within a cytoplasm(46). To suppress multiple nucleations, we had to combine fast and irreversible binding with a slow titration of Gag monomers into the volume, assembling in solution first before linking the assembled structures to the membrane via one lipid per monomer. The lattices in Fig 2 are a single connected continent, with imperfect edges and small regions with defects present in the tri-hexagonal lattice (Fig 2). These lattice topologies are in very good agreement with structures determined by cryoET, where the lattices also appear to be fully connected but defects and an imperfect edge are visible (10). Defects are expected in part because a sphere cannot be perfectly tiled by a hexagonal lattice, and due to the stochastic nature of assembly. In our simulations, we observed some defects in lattice ordering in the interior of the lattice, and incomplete hexamer order on the edges (Movie S1).

### B. A pair of Gag-Pol monomers will stochastically assemble adjacent to one another within the immature lattice

Although only 5% of the Gag monomers in our simulation are tagged as Gag-Pol (~125 out of ~2625 simulated proteins), we find it is extremely unlikely that a lattice will be assembled without a pair of them already adjacent (Fig 2B). This is due to the stochastic nature of the assembly and the fact that each monomer has 3 adjacent monomers, two via its hexamer interfaces and one via its dimer interface. Indeed, we find that even if we turn off any specific Gag-Pol to Gag-Pol binding interaction to mimic an unfavorable binding interaction during assembly (we set the binding rates to zero), we still find pairs of them adjacent, as they can be brought into proximity via their specific interactions with the Gag monomers (Fig 2B). This result indicates that to prevent early activation of the proteases, one cannot just rely on a low probability of Gag-Pol to Gag-Pol interaction, as the lattice is simply too densely packed. Instead, these Gag-Pol dimers would have to be actively inhibited from initiating protease activity, as any activation preceding budding is known to reduce infectivity(27).

### C. The Gag lattice disassembles with the weaker hexamer contacts of-5.62k_B_T

We perform all our simulations from the same starting structures, but with a range of hexamer strengths of-5.62 to −11.62k_B_T, and a slower (0.015μM^-1^s^-1^), medium (0.15) and faster (1.5) rate of binding for each ΔG_hex_. For the weakest hexamer contacts of −5.62k_B_T, we find that the lattice is not able to retain its single continental structure, and instead fragments into a distribution of much smaller lattices (Movie S2). Given a fixed Δ*G_hex_* we speed up the on- and off-rates and as expected, we see more rapid disintegration of the lattice structure. As we stabilize the lattice by increasing Δ*G_hex_*, *we* still see departure from the single continental structure due to unbinding of monomers and small complexes from the lattice edge (Fig 3). Hence, we see the emergence of a bimodal distribution of lattices, with a peak at the monomer/small oligomer end, and another peak containing the majority of the lattice in one large continent. The size of the large continent remains largest with increasing Δ*G_hex_* and with a slower rate, over the course of these ~17-20s simulations (Fig 3). Importantly, these dynamics occur in all our simulations and would not be possible if not for the incompleteness of the lattice. Specifically, the Gag contacts are dissociating not from the membrane but from each other, predominantly along the edge (Fig S3), at which point they can then diffuse along the membrane surface (Movie S1, S3). If the lattice were covering 100% of the surface, dissociation events would not allow Gags to diffuse away, and no dynamic remodeling would occur. From the sizes of the lattices present in the simulations, we can also report on the distribution of diffusion constants represented on the surface, as larger lattices diffuse more slowly. For the weaker lattices, the distribution is very broad, spanning 4 orders of magnitude, whereas for the most stable lattice there is primarily one very slowly diffusion time-scale, and a separate time-scale for the more faster moving oligomers (Fig S4).

**Figure 3.**
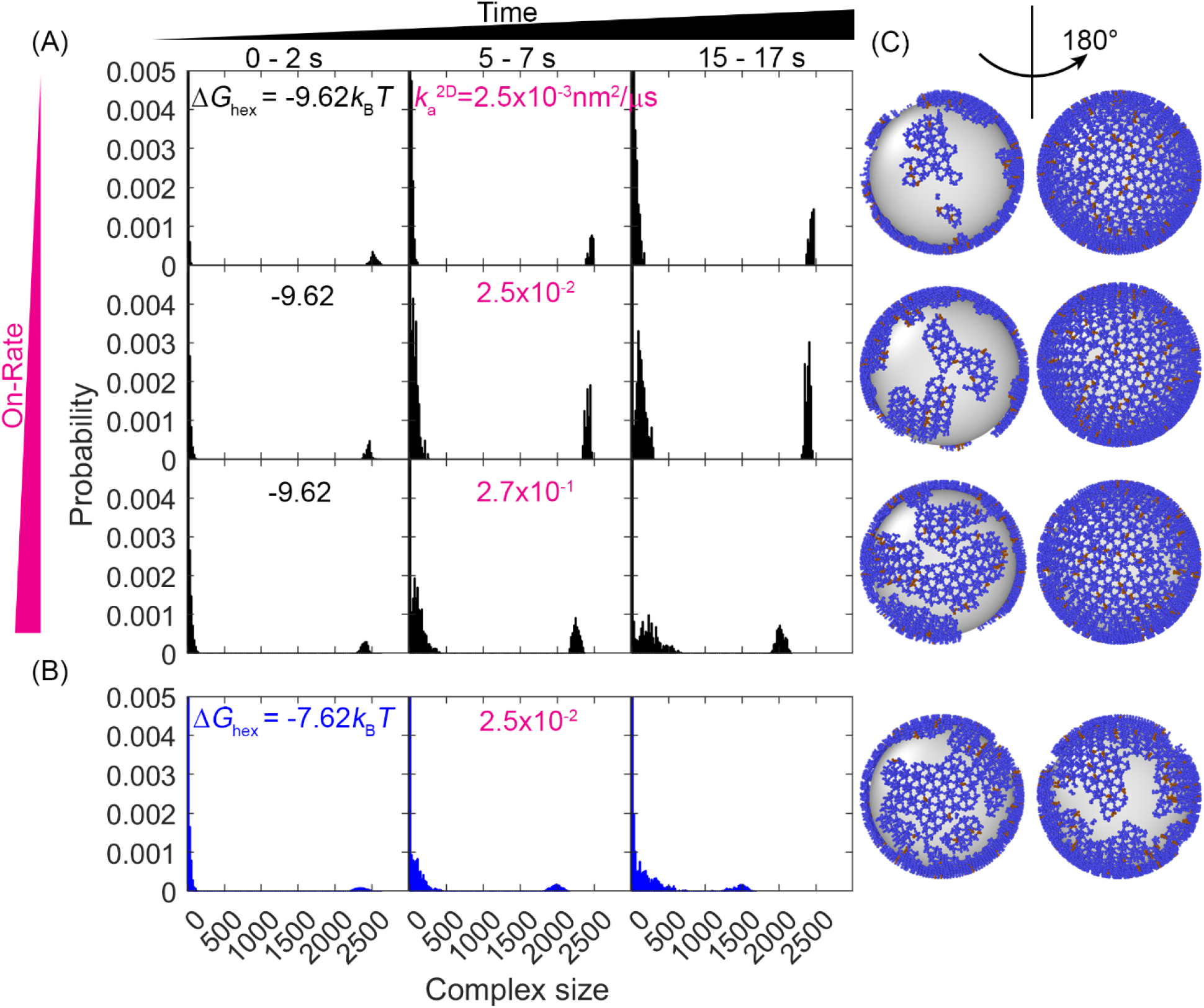
Evolution of the lattice size distribution at different reaction rates and hexamer interaction strengths. A) Along the x-axis are the numbers of monomers found in each lattice, which is largely bimodal for all systems: a population of small oligomers and one giant connected component. As time progresses (from left to right columns), the initial structure which was one giant connected component continues to fragment somewhat, indicating that the starting structure was not at equilibrium. As the on- and off-rates increase (from top to bottom) with a fixed Δ*G_hex_* = – 9.62*k_B_T*, the largest component shrinks, as shown by the peak denoting the large giant component shifting to the left, and the peak denoting the small oligomers shifting to the right. B) For a weaker hexamer free energy shown in the blue data (Δ*G_hex_* = – 7.62*k_B_T*), the lattice is breaking apart more rapidly and moving towards a more uniform distribution of lattice patch sizes as both peaks shift to the center. Note that we cutoff the y axis at 0.005 to make the peak at ~2500 visible. The plots extending the y-axis to 0.05 can be found in Fig S5. C) Representative structures at the later times (*t*=17s) for each case, illustrating the increased fragmentation as the rates accelerate, or as the hexamer contacts destabilize (lowest row).

### D. From the initialized lattice, Gag-Pol monomers can detach, diffuse, and reattach

A primary goal is to characterize the path and the timescale by which two Gag-Pol molecules could find one another given the single continental lattice they are embedded in at time zero. From our initial lattices, we showed in Fig 2 that at least one pair of Gag-Pol monomers are already in contact with one another, so we ignore those pairs to focus instead on spatially separated Gag-Pols. We can track the separation between all pairs of Gag-Pol monomers (Fig 4), and when this distance drops to the contact separation and persists, that indicates a binding event. We quantify the first passage time (FPT), or the time for the first pair to find one another in each simulated trajectory. We find that the incomplete structure of the Gag lattice supports a number of detachment events involving either a single Gag monomer or a small island of Gag lattice that can then diffuse along the surface and reattach at a new point (Movie S1). We quantify that the edge of the incomplete lattice is approximately 20 nm deep and contains ~250 Gag monomers. Of these monomers, 10% have only one link to the lattice, which offers the easiest path to disconnect, by breaking only a single bond (Fig S3). Ultimately, although only 5% are Gag-Pol monomers (N=125), we are nonetheless able to observe events that result in a pair of Gag-Pol monomers binding either through a dimer or hexamer contact, as we quantify further below.

**Figure 4.**
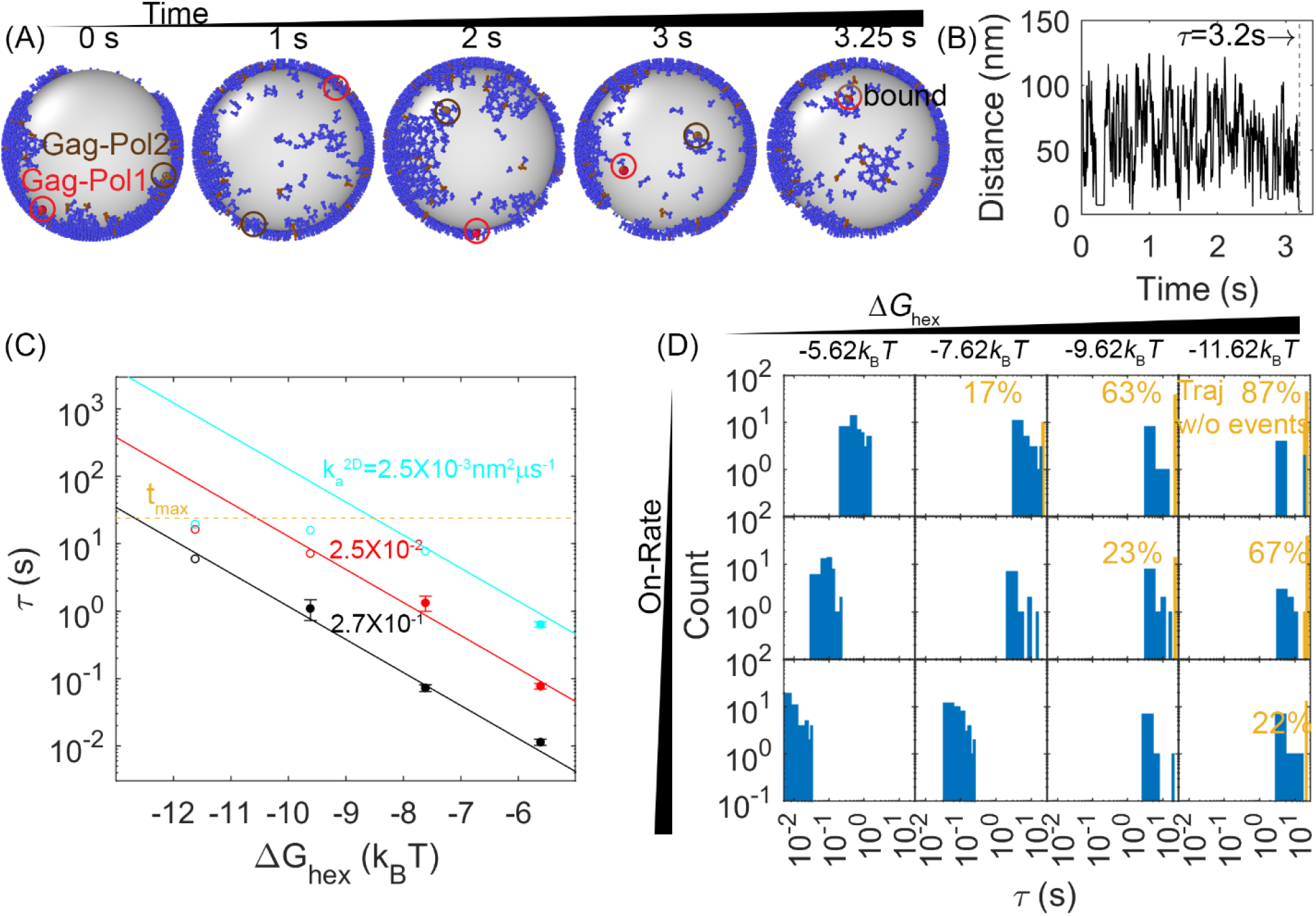
First-passage times for a pair of Gag-Pol monomers to search and bind to one another reveal a clear dependence on hexamer rates and free energies. A) An example from a simulation of how two Gag-Pols found each other. B) The distance between all Gag-Pol pairs can be monitored in time, with this trace corresponding to the simulation in (A). The distance fluctuates and drops to the binding radius σ at 3.2s, after which the two molecules remain bound. C) First passage times of Gag-Pol dimerization at different reaction rates and hexamer free energies Δ*G_hex_*. The yellow dashed line indicates the maximal length of the simulation traces. The filled-in circles report the MFPT for parameter sets where all traces produce a Gag-Pol dimerization event. The open circles report a lower bound on the MFPT, because some of the traces were not long enough to observe a Gag-Pol dimerization event. The solid lines are the fits to the FPTs from using Eq. 2, using only data points that had at least 75% of the trajectories produce dimerization events. D) The distributions of the first passage times at different reaction rates (each row) and hexamer free energy (each column). The yellow bars and yellow numbers report the percent of traces without any Gag-Pol dimerization event over the time simulated, so they are placed at the end of the simulated time. The first passage times slow as the reaction rates decreases and as the hexamer contacts become more stabilized.

### E. First-passage times for protease dimerization events are dependent on the free energy and binding rates of the hexamer contacts

In Fig 4, our results show how the first-passage time for two Gag-Pols to dimerize with one another is dependent on both unbinding rates and binding rates, as both events are required to bring two Gag-Pol together. We observe two intuitive trends. One is that for a given free energy Δ*G_hex_*, a faster association (and thus faster dissociation) rate results in faster dimerization events between the Gag-Pol monomers. The second trend is that as the lattice free energy stabilizes, dimerization events are slowed due to the slower dissociation times, despite having the same on-rates (Movie S3). These timescales are thus consistent with the dimerization events requiring at least one of the monomers to dissociate from the lattice, and then rebind at a new location containing a Gag-Pol. We report the association rates as their 2D values, because binding is occurring while the proteins are affixed to the 2D membrane surface, as unbinding from the surface is rare (Methods). The corresponding 3D rates are representative of slow to moderately fast rates of protein-protein association (1.5×10^4^ −1.5×10M^-1^S^1^), where they are converted to 2D values via a molecular length-scale h=10nm (Methods). Activation requires explicitly that the 5% of monomers carrying the proteases to be involved. Hence, additional unbinding and rebinding events will occur that are not ‘activating’ because they involve a Gag monomer without a protease. Gag-Pol molecules can also unbind and rebind multiple times before successfully finding another Gag-Pol.

### F. Mean first-passage times can be well-approximated and predicted as a function of Δ*G*_hex_ and binding rate *k*_a_

For the models with weaker and/or faster interactions, our simulations were long enough that all trajectories of that model (*N*_traj_~60) resulted in a Gag-Pol dimerization event. We could thus construct the full first-passage time distribution and reliably calculate the mean first passage times, *τ_MFPT_*. The MFPTs displayed a remarkably clear functional dependence on both the Δ*G*_hex_ and *k*_a_ values as shown in Fig 4, for models where the sampling was complete. In contrast, for the most stable lattices with the slowest rates, the majority of the trajectories had not yet produced a dimerization event (Fig 4D—yellow bars), and thus the simulations do not report on the true MFPT. We therefore derived a formula to fit the completed MFPT values, first by analogy to a MFPT model for bimolecular association, with an inverse dependence on *k*_a_ (see, e.g. (43)). Second, we empirically find that the *τ_MFPT_* also has a power-law dependence on the *K*_D_, *τ_MFPT_* ∝ *K_D_^γ^*, or equivalently, 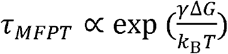. Our phenomenological formula thus had two fit parameters, the power-law exponent *γ* and a constant pre-factor (see Methods). After fitting, we find the approximate relationship:

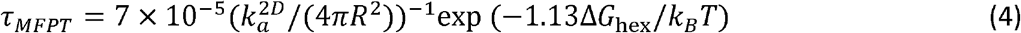

Where 7×10^-5^ is a dimensionless fit parameter, *R* is the radius of the sphere, and our convention has Δ*G_hex_* < 0. We see excellent agreement between our formula and our data (Fig 4C), which allows us to extrapolate the remaining models beyond the maximum simulation time of 20 seconds.

### G. For our models, activation events occur in less than a few minutes

Importantly, in the models we have studied, the activation of a dimer can occur in well under a minute up through several minutes (Fig 4). For hexamer stabilities of −5.62 and −7.62k_B_T, all rates support dimerization events at less than 10s. For the more stable lattices of −9.62 and −11.62 k_B_T, only the medium and fast rates ensure a MFPT that is less than or comparable to (~50s): an event occurring within 100s of the start. Using our Eq. 4, we can determine that for a moderate rate of 2.5×10^-2^nm^2^μs, a Δ*G*_hex_ more stable than −12k_B_T will be slower than 100 seconds. For the slowest rate of 2.5×10^-3^nm^2^μs^-1^, anything more stable than −9.4k_B_T will be slower than 100 seconds. Our most stable lattices at the slowest rates take 10 minutes on average for an activation event. Our results thus quantify and predict how the kinetics and the stability of the lattice must be tuned to allow sufficiently fast dimerization events involving the 5% of Gag-Pol molecules carrying polymerases.

### G. Lower lattice coverage does not dramatically change the first-passage times

When comparing lattices with 66% coverage vs 33% coverage, we see in some cases a minor slow-down in dimerization times, but the MFPT is overall much less sensitive than it is to the binding rates. With 33% coverage, the edge of the lattice does have a comparable size to the 66% lattice, but the ‘bulk’ interior is smaller, with more free space required to diffuse to a partner. However, most significantly, the concentration of Gag is smaller, and now with only 66 Gag-Pols (vs 125) present in the lattice, we see in some cases an increase in the time it takes for a pair to find one another (Fig 5A).

**Figure 5.**
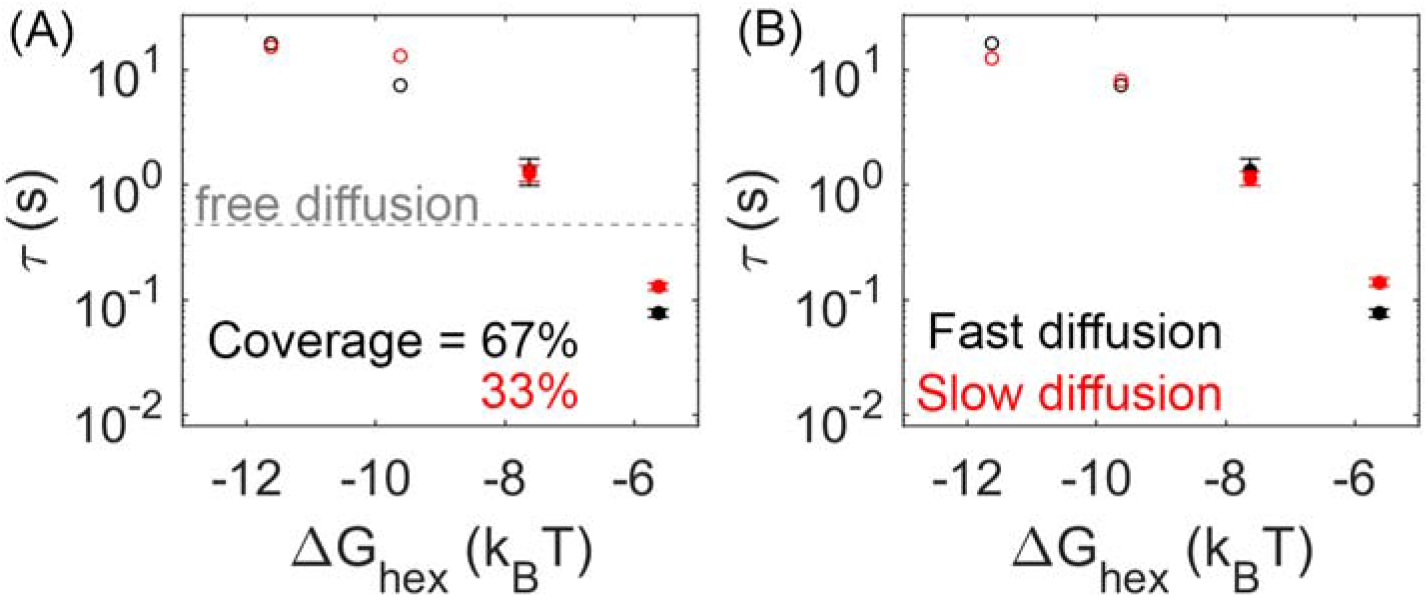
Lower lattice coverage or slower diffusion does not dramatically change the mean first-passage time. A) First-passage time at different lattice coverages (67% and 33%). Closed circles are cases where Gag-Pol dimerization occurs in 100% of trajectories, while open circles are cases where Gag-Pol dimerization occurs in < 100%. The gray line is the first-passage time when two free Gag-Pols diffusing on an empty spherical surface bind to one another. For the weakest lattice, rebinding is actually faster than diffusional encounter times between a dilute pair. B) Mean first-passage time at different diffusion constants of the lipid. A monomer of Gag on the membrane diffuses at 0.2 um^2^/s (black data), and diffuses slower as it grows in size consistent with Einstein-Stokes (Methods). We also simulated the system where diffusion of all species was slowed by a factor of 10 (red data).

### H. First passage times are not sensitive to diffusion or dimerization strengths

We tested two strengths for the dimerization free energy of ΔG_dimer_ −11.62 k_B_T and −13.62 k_B_T, comparable to experimentally measured values of dimerization in solution(39). The MFPT were overall relatively similar across both values, indicating that the timescales do not show the same sensitivity to Δ*G*_dimer_ as Δ*G*_hex_ (Fig S6). This likely emerges because the dimer is more stable in most cases than the hexamer. Thus the hexamer unbinding events are more frequent and more likely to directly provoke the first activation events. Further, the hexamer has two binding sites, so more contacts in the lattice, and because hexamers nucleate stable cycles needed for higher order assembly, the frequency of hexamers vs incomplete hexamers are significantly more sensitive to Δ*G* than a single dimer bond. This result shows that breaking and formation of the hexamer contacts is important in driving Gag-Pol dimerization events, given that the MFPT shows clear sensitivity to these rates.

We similarly found minimal dependence of the MFPT on the diffusion constant. With a 10-fold slower diffusion constant for all membrane-bound Gag monomers, which effectively slows all lattices down by 10-fold, the MFPT were not significantly slower (Fig 5b). This is not surprising given that the diffusional search along the membrane to find a new partner is not ultimately the rate-limiting step in the association process. The rates we report are the intrinsic rates, which control binding upon collision, whereas the effective macroscopic rates are dependent as well on diffusion. Faster intrinsic rates are more diffusion-limited and produce binding that is more sensitive to diffusion (38). However, given the small dimensions of the virion, traveling ~70nm for example (the radius) takes on the order of milliseconds for monomers and small oligomers of Gag. Rough estimates of delay times for a binding event, using 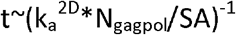 indicate that even for the fastest binding, it is on the order of a few milliseconds. The slowest timescale given all the rates is for the stable lattice, where dissociation has a timescale of ~7 seconds. Altogether, these timescales of individual steps show that the observed MFPT are not merely controlled by the slowest single events, but by the need for multiple attempts of un- and re-binding to ensure a pair of Gag-Pols find one another. Our results further illustrate how the crowding due to the lattice on the surface can actually accelerate rebinding events compared to a freely diffusing pair when the lattice is unstable (Fig 5a), whereas for stronger Gag contacts the lattice will dramatically slow rebinding.

### I. Biochemical measurements of Gag mobility in VLPs agree with our moderately stable lattices

We find that the dynamics of our simulated lattices agrees with experimental measurements of binding within the lattice for parameters that exclude the most stable, slowest regimes. Experimental measurements within the Gag lattice of budded virus-like particles (VLP) tracked the biochemical formation of a Gag dimer involving a population of Gag molecules tagged with a SNAP-tag (10-40%) and the same fraction of Gag molecules tagged with a HALO-tag(32). A covalently linked dimer was formed through addition of a HAXS8 linker at time zero, with one linker forming an irreversible bridge between a HALO and SNAP protein. Formation of this covalently linked dimer was quantified to reveal an initial rapid formation of dimers, followed by an increasing slower growth that reaches 42% dimer pairs formed for the 10% tagged populations. In our simulations, we thus performed a comparable ‘experiment’ given our trajectories (see Methods). We tracked the encounter between two populations of our Gag molecules that had been randomly tagged as either 10% Halo or 10% SNAP (Fig 6). We similarly found that the majority of the dimers formed rapidly, because they were already adjacent in the lattice when the covalent linker was introduced. The dynamics of the lattice then allowed a slow growth in additional dimers (Fig 6a). We calculated the fraction bound over the course of our 20 second simulations and used a simple extrapolation to define an upper bound on the number formed at 3 minutes (see Methods). For the least stable lattices, the upper bound is close to all dimers formed, over all 3 association rates, which is much higher than observed experimentally. For the most stable lattices in contrast, even when our model assumes maximal efficiency of the covalent linker, we extrapolate to an upper bound that is less than 42% dimers formed experimentally (Fig 6b). These results thus indicate that these lattices are too stabilized to support the dynamics observed in the Gag VLPs.

**Fig 6.**
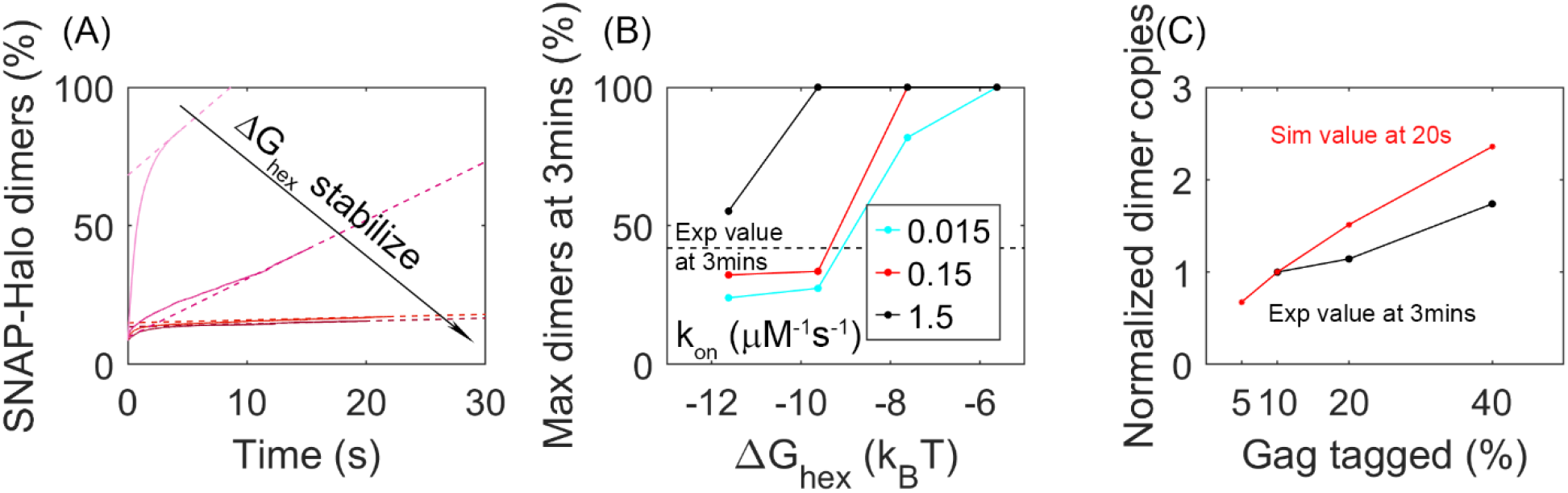
Analysis of our simulations mimics experimental biochemical measurements of Gag dimerization as a function of time, agreeing for moderately stable lattices. **A)** Percent of tagged Gag molecules that have formed a dimer (involving a SNAP-tag and HALO-tag plus linker) as a function of time. Here, 10% of Gag monomers were initially tagged either HALO or SNAP. As the hexamer stability Δ*G*_hex_ increases, the dimerization yield dramatically slows. The dashed lines are linear fits of the last 1s of the curves. Results averaged over all 60 traces per parameter set. B) Yield of dimers formed at 3 minutes estimated via simple linear extrapolation. Our results represent an upper-bound. Dashed black line is the experimental measurement of the dimer formation at 3 mins given 10% tagged populations. Fast (black), moderate (red) and slower (cyan) rate constants. With the most stable lattices and slowest rates, dimer yield is too low compared to experiment. C) Dimer yield as we increase the population of initially tagged Gag monomers from 5% to 40%. Red is simulated yield at 20s (to avoid extrapolation assumptions), and black is experimental yield at 3mins. We normalize the yield by the value at tagged Gag=10%, given the different time points used.

We also verified that these simulations are consistent with the trends expected as the population of tagged Gag monomers is increased. Indeed, we find, similar to experiment, that as a larger fraction of Gag monomers have tags, corresponding to a higher concentration of binding partners, we see a larger fraction of dimers being formed (Fig 6C).

### J. Large-scale and heterogeneous lattice dynamics are visible in auto-correlation functions, and are qualitatively similar to iPALM experiments

We quantify the dynamics of the Gag lattice on fixed viewpoints on the spherical surface using number auto-correlation functions (ACFs), which report on correlations of collective motion that can emerge due to heterogeneity within the lattice (Fig 7) (Methods). We expect this heterogeneity due to our lattices all exhibiting a large component and smaller oligomers (Fig 2). We find that as the lattice becomes more stable and the bimodal separation of lattice sizes becomes more pronounced, the measured correlations in Gag copy numbers per quadrant increase in amplitude and slow in timescales, and that these dynamics are sensitive to slowing diffusion (Fig S7). All the ACFs will eventually asymptote to 1 at long delay times as the copy numbers become independent (Fig 7a).

**Fig 7.**
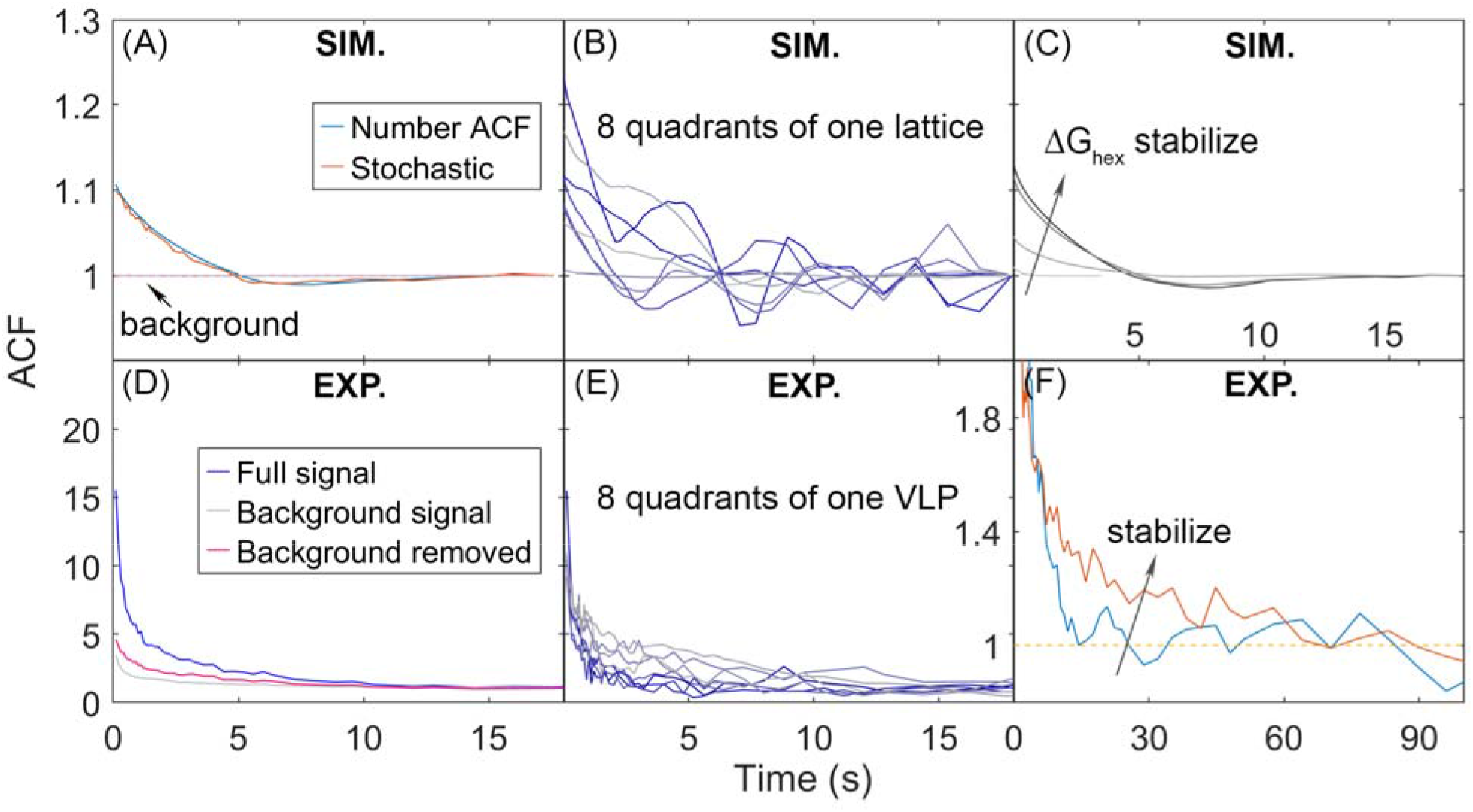
Auto-correlation functions (ACF) of lattice dynamics from simulation and experiment show qualitatively similar trends. A) Number ACF of the simulations calculated directly from the copy numbers of Gag monomers shown in blue. Averaged over all 8 quadrants over all 60 traces for one parameter set. Using the stochastic localization method shows excellent agreement (Orange). Dashed lines are the background signal, which is 1 as expected (bleaching of the Gag monomers causes limited drops in total copies across 20s), as the total copy numbers across the membrane surface do not change. We note that the ACF values at our longest delays (i.e. *τ*>~10s) are not statistically robust, because of the limited number of frames separated by these timescales. B) ACF of each of the 8 quadrants of one simulated lattice. C) As the lattice is stabilized by increasing Δ*G*_hex_, the ACF shows higher amplitude correlations that decay to 1 at longer times D) ACF from stochastic localization experiments on Gag VLPs. The blue curve is the average signal over all 8 quadrants over 11 VLPs. The gray is the background signal for the ACF of the total copy numbers across the surface, then averaged over all VLPs. The red line is the ACF signal after dividing out the background. E) ACF of 8 quadrants of one experimental VLP. F) The ACF from VLPs that have been stabilized with a fixative (orange curve), show the same trend as the stabilized lattices from simulation. The y-axis has been zoomed in to demonstrate the shift.

In Fig 7, we show how the ACFs calculated from simulation show similar behavior to the ACFs measured from experiment. Each of the 8 quadrants of the spherical surface (4 on the top hemisphere, 4 on the bottom hemisphere) displays heterogeneity in the amplitude of correlations because some quadrants contain large lattice fragments, and others contain mostly empty space (Fig 7b). The same trend is observed in a single VLP measured experimentally (Fig 7e). Our simulations further show that as the lattice is stabilized, the ACF increases in amplitude and decays more slowly, which is qualitatively the same as is observed in iPALM experiments of Gag lattices in budded VLPs that have been stabilized with a fixative(25) (Fig 7c, 7f). We cannot quantitively compare the ACFs, as the experiments produced ACFs with much higher amplitudes of correlations, and even with the background correlations divided out (Fig 7d), the experimental signal contained additional sources of correlation likely due to measurement noise. However, we were able to use our simulations to illustrate how sources of measurement noise in stochastic localization experiments can produce increased correlations beyond the background. We specifically find that short-term blinking of the fluorophore does not appreciably change the ACF (Fig S8). We do see increased amplitude of correlations in the ACF if we introduce a distribution of activation probabilities for the fluorophores, mimicking the fact that the populations initially activated may have a higher probability of activation than those appearing at later times (Fig S8). Further, if we assume that the lattice is not perfectly centered with respect to the activating laser pulses, then Gag monomers that are initially ‘dark’ can diffuse into view and then have a probability of being activated (Fig S8). This increases fluctuations in both the background signal and the signal from the separate quadrants. Thus, the simulations improve interpretation of the experiment, and more vividly bring to life the lattice dynamics and heterogeneity.

## DISCUSSION

Our simulations of Gag lattices across a range of interaction strengths and rate constants all demonstrate how the incomplete lattice supports dynamic unbinding, diffusion, and rebinding of Gag along its fragmented edge. These dynamics and the resultant accessibility of Gag molecules along the edge of the lattice are important for activation of proteases via dimerization, and ultimately the maturation of the virion from this spherical lattice shell to the mature capsid. By measuring the first-passage time for dimer formation between a pair of Gag-Pol monomers, we show that dimerization can proceed in less than a few minutes, and for less stable lattices much faster, despite the embedding of these molecules within the lattice. By comparison with experimental measurements of lattice structure (via cryoET) and lattice dynamics (via biochemistry and iPALM), we conclude that the stability of the hexamer contacts should be in the range of — 12*k_B_T* < Δ*G*_hex_ < – 8*k_B_T* for binding rates that are slower than 10^5^M^-1^S^-1^. If the binding rates are faster, than the free energy could be further stabilized, as the dimerization events would still be fast enough to be consistent with the biochemical measurements. If the lattice is less stable than – 6*k_B_T*, we found that the large-scale structure of the lattice is not maintained even within seconds, which is not consistent with structural measurements, and between – 6*k_B_T* to — 8*k_B_T*, the dimerization is likely too fast relative to the biochemical measurements. These hexamer-hexamer contact strengths report the stabilities that would be expected given the presence of co-factors, as without co-factors the lattice does not assemble at all (47, 48), and co-factors are present in all experiments used for comparison. Our simulations also demonstrate that during assembly, the fraction of Gag-Pol monomers, while only 5%, is still too high to prevent stochastic dimerization events between them. This means that preventing early activation, which can result in loss of proteases from the virion(49) and significant reduction in virion formation (27), requires active suppression of the interaction between adjacent Gag-Pol monomers.

Our model explicitly accounts for the crowding effects of localizing the lattice to a small, 2D surface, ensuring that excluded volume is maintained between all monomers. However, we do not explicitly include the genomic RNA that would be packaged within the immature virion and attached to the Gag lattice (through non-competing binding sites), or other important assembly co-factors such as IP6. It is known that binding to RNA (50, 51), membrane, and other co-factors is important in stabilizing the lattice for assembly (18, 21, 22, 39, 52). Thus we do implicitly account for their influence in stabilizing hexamer free energies because the monomers support assembly. Furthermore, our Gag monomers and oligomers diffuse along the membrane surface, not through the interior of the budded virion where the RNA would be packaged. When our lattice coverage changes from 33% to 66%, for example, we see only small changes in our mean-first passage times which primarily reflect the increase in total Gag-Pol monomers available. However, the attachment of the Gag lattice to a large RNA polymer of 9600 nucleotides (~3μm) could change the mobility of the Gag monomers following their detachment from the lattice. Proteins can still unbind and diffuse when bound to a polymer like RNA(31, 53), but the effective rates could slow, and the distance that a monomer typically travels can be limited by the fluctuations of the attached RNA polymer. Hence rebinding may be more restricted to shorter excursions from the start point. We note the Gag VLPs contain smaller RNA polymers and not the full gRNA, so the model is in that way more consistent with the dynamics of a VLP. Somewhat remarkably, the Gag monomers within the virion are able to reassemble around the gRNA to form the mature conical capsid (following cleavage)(41), which indicates there is a clear capacity for diffusion driven remodeling. This mature lattice is also subsequently disassembled(54), and the principles of our model here indicate how destabilization of hexamer contacts could help promote disassembly.

Our models here contain pairwise interactions, and cooperativity enters only in that the formation of a completed cycle (whether a hexamer or a higher-order cycle of multiple hexamers) is significantly more stable, because it requires two bond breaking events. However, coordination of the hexamer by IP6 can produce conformational changes(48) or kinetic effects(30) that could change the stability of hexamer contacts between say, a dimer vs a 5-mer. We did not include this additional cooperativity to keep the model as simple as possible; we expect that added cooperativity in hexamer formation would change the prefactors in the quantitative relationship we predict between the hexamer free energies and the first-passage times, as intermediates would be biased away from smaller fragments. However, because the lattice would inevitably still have the ‘dangling’ edges and partial hexamers observed experimentally(10), we would still see dissociation, diffusion, and rebinding events. Lastly, other proteins are packaged into HIV-1 virions, including curvature inducers(55), and similar to RNA, additional protein interactions could shift the Gag unbinding kinetics. Ultimately, however, our models clearly show that despite the significant amount of protein-protein contacts and ordered structure within the membrane attached Gag lattice, there is nonetheless enough disorder along the incomplete edge to support multiple unbinding and rebinding events over the seconds to minutes time-scale (Movie S1, S3).

Overall, the model and simulations here reveal a level of detailed Gag dynamics coupled to structural changes that are inaccessible to any single experiment but can nonetheless be compared to a range of experimental observables, as we have done here. Although diffusion does influence the collective dynamics of the lattice, for example, we find it does not significantly influence activation rates, as those are limited by binding and unbinding events rather than mobility. By defining a formula that allows us to extrapolate our model to other rates and free energies, we can predict how mutations that would change the strength or kinetics of the hexamer contacts would impact the time-scales of the initial protease dimerization event. Mean first-passage times can be predicted from theory in surprisingly complex geometries (56), but for the immature lattice, the problem was intractable without using simulation data due to the ability of Gag-Pol to rebind or ‘stick’ back onto the lattice through multiple contacts before successful dimerization encounters. More generally, modeling stages of viral assembly has been critical for establishing the regimes of energetic and kinetic parameters that distinguish successful assembly from malformed or kinetically trapped intermediates, such as in viral capsid assembly (57–60). Computational models of self-assembly can be used to assess how additional complexity encountered *in vivo*, such as macromolecular co-factors (30, 31, 61), crowding (59, 62), and changes to membrane-to-surface geometry (24), could help to promote or suppress assembly relative to *in vitro* conditions. Our reaction-diffusion model developed here provides an open-source and extensible resource (34) to study preceding and following steps in the Gag assembly pathway with the addition of co-factors. With rates and energies that match biochemical measurements, the model can act as a bridge between *in vitro* and *in vivo* studies of retroviral assembly and budding, and a tool to predict assembly conditions that disrupt progression of infectious virions.

## AUTHOR CONTRIBUTIONS

MEJ and SS designed the research. SG and IS performed simulations. SG and MEJ generated analytic tools. SG analyzed data and generated figures. SG and MEJ wrote the initial manuscript. All authors approved the final manuscript.

## ACKNOWLEDGEMENTS

M.E.J. gratefully acknowledges funding from an NSF CAREER Award 1753174. The funders had no role in study design, data collection and analysis, decision to publish, or preparation of the manuscript. We acknowledge use of the ARCH supercomputer Rockfish at Johns Hopkins, with support from NSF MRI 1920103 and the XSEDE supercomputer Stampede2 through XRAC MCB150059. Contributions from I.S and S.S. were supported by NIH R01 AI150474.

## SUPPLEMENTAL MOVIE LEGENDS

**Movie SI:** Lattice dynamics for moderately stable dynamics that are consistent with structural and biochemical experiments. Δ*G_hex_* = −9.62*k_B_T*. *k*_a_^2D^ (nm^2^/μm) = 2.5×10^-2^; gap between each frame: 10ms; overall time length: 20s.

**Movie S2:** Lattice dynamics for an unstable lattice. Δ*G_hex_* —S.62*k_B_T*. *k*_a_^2D^ (nm^2^/μS) = 2.5×10^-2^; gap between each frame: 10ms; overall time length: 4.5s.

**Movie S3:** Lattice dynamics for a more highly stable lattice with slower binding kinetics. Δ*G_hex_* –-11.62*k_B_T*. *k*_a_^2D^ (nm^2^,μS) = 2.5×10^-3^. gap between each frame: 10ms; overall time length: 20s.

## SUPPORTING REFERENCES

References (XX-XX) appear in the Supporting material.

