## Supplemental Information for "Defects in the HIV immature lattice support essential lattice remodeling within budded virions"

Supporting Information

### SUPPLEMENTAL METHODS

**Orientational Angles between Gag-Gag contacts to ensure the spherical lattice assembly:** We derived the Gag monomer structure and the angles needed to orient a pair of bound Gags directly from the PDB structure 5L93.pdb. One exception is the MA domain, which binds the membrane and is not present in the PDB structure. We chose the position of the MA domain such that the lattice would assemble properly on the membrane surface, with each monomer binding along the normal to the surface. The binding sites positions, binding radius, and binding angles are listed in Table S1.

Table S1. The binding sites positions, binding radius, and binding angles of the coarse model in Fig 1. The position of center-of-mass (COM) is [0,0,0] nm. $\theta_1, \theta_2$ are the angles between the binding radius and the site-to-COM vector. $\varphi_1, \varphi_2$ are dihedral angles that orient a second axis of each molecule relative to the binding radius. $\omega$ is a dihedral angle between the site-to COM vectors around the binding radius.

| | |
|---|---|
| MA domain (nm) | [-0.08, -0.004, 1.998] |
| Dimerization site (nm) | [-1.01, -1.244, -1.175] |
| Hexamerization site 1 (nm) | [1.277, -1.009, -2.087] |
| Hexamerization site 2 (nm) | [1.095, 0.313, -2.12] |
| MA-Membrane binding radius (nm) | 1 |
| Dimerization binding radius (nm) | 2.21 |
| Hexamerization binding radius (nm) | 0.42 |
| Dimerization binding angles $[\theta_1, \theta_2, \varphi_1, \varphi_2, \omega]$ rad | [2.168, 2.168, -2.005, -2.005, -1.723] |
| Hexamerization binding angles $[\theta_1, \theta_2, \varphi_1, \varphi_2, \omega]$ rad | [2.21, 1.459, 2.443, 1.55, 0.879] |
| MA-membrane binding angles $[\theta_1, \theta_2, \varphi_1, \varphi_2, \omega]$ rad | $[\pi, \pi$, -,-,-] |

***Observed association rates:*** For the kinetics of association of the Gag monomers in 2D, we validated our model parameters against analytical or numerical solutions for simpler systems. To test dimerization kinetics, we turn off the hexamer interactions and initialize all monomer irreversibly on the membrane. We found that the rates that we assigned via the input files, for instance $k_a^{2D}$=0.02 $nm^2/\mu s$, when simulated, produced kinetics with an apparent rate that was 4x slower. This same factor of 4 slow-down was observed for input rates of 0.2 and 2 $nm^2/\mu s$. The dissociation kinetics are unaffected. We found that the reason for the slower observed kinetics, and thus the slower apparent association rate, is because we reject association events that cause a reorientation of the monomers into the bound state that is larger than our specified threshold. Our threshold is controlled by a scale-factor (scaleMaxDisplace) that scales the expected displacement of each monomer prior to association. This scale-factor was set low enough (to 10) for the slowly diffusing and non-rotating 2D monomers that events were rejected. We therefore report throughout the paper the rates that describe the actual observed kinetics, and are thus 4x lower than the values we specified in our input files. This comparison as shown in Fig S1 then agrees very well between simulation and theory. We performed the same validation for the Gag monomers to form hexamers, with the dimer interaction turned off but still excluding volume. Here again we found the same 4x slow-down in association kinetics relative to the assigned rates in the input file. Therefore, we always report these apparent rates as shown in Fig S2 that accurately describe the kinetics.

### ***Simulations of biochemical linkage between SNAP-tagged Gag and HALO-tagged Gag via the covalent linker HAXS8***

For our simulations of dimerization between a SNAP and HALO-tagged Gag monomers, we chose the 3nm cutoff distance because it is comparable to the molecular length-scales of the two protein tags with the linker between them(44). Any encounter at this length was considered a binding event, so the rate was maximally strong, and all events were treated as irreversible, which is consistent with the covalent bond being formed by the linker to both tags. That meant that neither of the two monomers would be able to participate in any other events. This model assumes that the arrival of the linker to the inside of the virion is relatively rapid. The permeability coefficient of the linker when exposed to the membrane enclosed Gag lattice is approximately 0.0004nm/us [1], and assuming a membrane thickness of ~5nm, the diffusion across the membrane occurs at ~0.002nm2/us. To test the role of linker permeability, we solved the diffusion equation for a 1uM concentration of linker molecules diffusing into of a sphere of radius R=67nm, which mimics the experiments. Within 100ms, the concentration of the linker at 60nm (close to the Gag-tagged end) has already reached 0.8uM. Hence although there is some delay following addition of the linker, it is much less than the time (20s – 3 minutes) over which most of the dimerization occurs. The model also assumes that the linker does not saturate all HALO and SNAP molecules independently, which would prevent any dimers forming. From the literature, we found that the rates of binding of SNAP and HALO to the linker HAXS8 are $3x10^4$ and $3x10^6$ respectively[1]. The linker HAXS8 binds to both proteins specifically, and not to other linkers. Therefore, if all HALO and SNAP are bound to linkers first, then no dimers can form. We solved a system of ODES for binding of HALO and SNAP to a linker given the rates listed above. The HALO and SNAP concentrations were controlled by the size of the virions with 250 of each present (10%), and the linker concentration was 1uM, which was found experimentally to ensure high dimerization success. The linker binds more rapidly to HALO, but there is still plenty of time for the HALO-linked molecules to bind to a free SNAP before all the sites are occupied by linkers, as the copy numbers of linkers in the volume are actually quite low (on average 1 at a time). In particular, if the HALO and SNAP tags are adjacent in the lattice, the SNAP is much more likely to bind the adjacent HALO-linker than a free copy. So it is reasonable to assume that a large fraction of successfully linked dimers can form. This also contributes to the model predictions being an upper bound on total dimers formed.

***Background removal from the ACFs:***

For the ACFs, we aim to measure how concentrations of monomers in a fraction of the virus surface are correlated in time. Across the 8 quadrants, we then measure the copies of monomer per quadrant: $n_1(t)$, $n_2(t)$, ... $n_8(t)$. The total copies are then $N(t)= n_1(t)+n_2(t)+ ... +n_8(t)$. The ACF for any single quadrant is given by $ACF(\tau) = \frac{\langle n_i(t+\tau)n_i(t)\rangle}{\langle n_i(t+\tau)\rangle\langle n_i(t)\rangle}$. For the total surface, $ACF(\tau) = \frac{\langle N(t+\tau)N(t)\rangle}{\langle N(t+\tau)\rangle\langle N(t)\rangle}$, which we denote as the background signal. From the simulation, the background ACF is 1 at all times, as total copies do not change. For the localization method, there is a drop in total copies due to 'bleaching' of molecules we localize, but it does not noticeably affect the background ACF. In the experimental measurement, the background ACF does exhibit correlations, indicating that the total copy numbers detected across the surface are varying in time. To remove this effect of total copy number variations, and instead focus on the local fluctuations in concentrations per quadrant, we would like to report a corrected $ACF_{corr}(\tau) = \frac{\langle n_i(t+\tau)/N(t+\tau)n_i(t)/N(t)\rangle}{\langle n_i(t+\tau)/N(t+\tau)\rangle\langle n_i(t)/N(t)\rangle}$. However, we do not have access to the relative concentrations $n_i(t)/N(t)$ at each time-step. A reasonable approximation is to assume that we can separate the average behavior of $n_i(t)$ and $N(t)$, such that $ACF_{corr}(\tau) \approx \frac{\langle n_i(t+\tau)n_i(t)\rangle/\langle N(t+\tau)N(t)\rangle}{\langle n_i(t+\tau)\rangle\langle n_i(t)\rangle/\langle N(t+\tau)\rangle\langle N(t)\rangle}$, which is equivalent to dividing out the background ACF from the signal of each quadrant. This background corrected ACF reproduces the exact ACF when the total copy numbers are constant.

***Modifications of the stochastic ACF calculation to introduce additional sources of correlation:*** For the original stochastic ACF calculation method, we set the probability $p_{act}$ = 0.6. We found that lowering this activation probability did not change the amplitude or timescales of the ACF, it simply made the signal more noisy. To introduce additional sources of correlation, one property we tested was to add fluorophore blinking. Each molecule can be localized up to 3 times within the 10 frames since its first localization, based on experimental characterization of the Dendra fluorophore[2]. Each fluorophore is on average observed twice, but always within 1s of its first appearance. A second property we tested, is we set the localization probability to decay with time: $p_{act}$ = 0.6exp(-t/10) to mimic that not all fluorophores are activated evenly, and the ones observed early on were more likely to be activated then those observed later. A third property we tested is to introduce 'dark' populations using two methods to mimic a limited activation region of the laser. 1) Only monomers in the top 4 quadrants are set to visible to the laser activation. Thus any molecules in the lower half are effectively 'dark' until they diffuse into the top half of the surface. We thus analyze only 4 quadrants. 2) The lattice is shifted relative to the laser focus to a distance 20nm along the x-axis and z-axis. Then, only the molecules whose distance to the origin is less than $R_{sphere}$=67nm are visible, and an asymmetric dark region exists. Molecules in the dark region cannot be activated, but if they diffuse into the visible region then they can be activated. Both these methods introduce correlations in the background signal, due to total copy number fluctuations, which also increases the amplitude of correlations in the signal.

**SUPPLEMENTAL FIGURES**

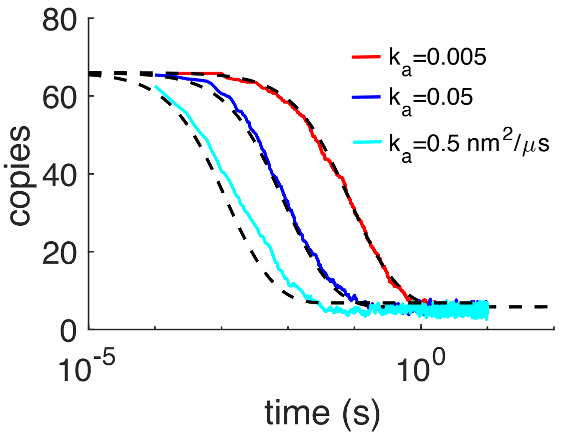


**Figure S1. Kinetics of dimer formation between Gag monomers is consistent with theory**. In these simulations, only the homo-dimer contacts could form, and the hexamer sites were turned off. Thus, the kinetics of reversible dimerization from NERDSS simulations (colored lines) can be compared with the non-spatial rate-equation solution (black dashed). Here we initialized all monomers to be on the membrane surface irreversibly, so the binding is purely in 2D. For each model, 5-10 trajectories were collected and averaged. As the microscopic association rate $k_a$ was increased, we also increased the dissociation rate $k_b$ the same amount, so that the free energy was fixed for all simulations at -11.9$k_BT$. The apparent 3D rates for the slowest system were $k_a^{3D}$=0.025 nm$^3$/$\mu$s and $k_b$=0.1s$^{-1}$. The length-scale to convert from 3D rates to 2D rates was set here to $h$=5nm, hence the slowest 2D rate is $k_a^{2D}$=0.005 nm$^2$/$\mu$s. The initial copy numbers were 66 on the same membrane surface as the full assembly simulations (a sphere with $R$=67nm). Consistent with the full system simulations, diffusion of each monomer on the membrane is set to 0.2 $nm^2/\mu s$ and the binding radius for the dimer interaction is $\sigma$=2.21nm. For the analytical solution in black dashed, we input the corresponding macroscopic 2D rates $k_{on}^{2D}$ and $k_{off}^{2D}$ . The macroscopic on-rate $k_{on}^{2D} \leq k_a^{2D}$ due to its dependence on diffusion constants and the system size[3]. We note that the agreement is not perfect between the reaction-diffusion simulations and the non-spatial solution. Although the kinetics are not expected to be identical in 2D due to sensitivity to spatial fluctuations, there is some disagreement because in the RD simulations, some of the association events were rejected if the monomers were not aligned closely enough to their target bound state, which causes a relative slow-down in the association rates. See SI Methods for more details.

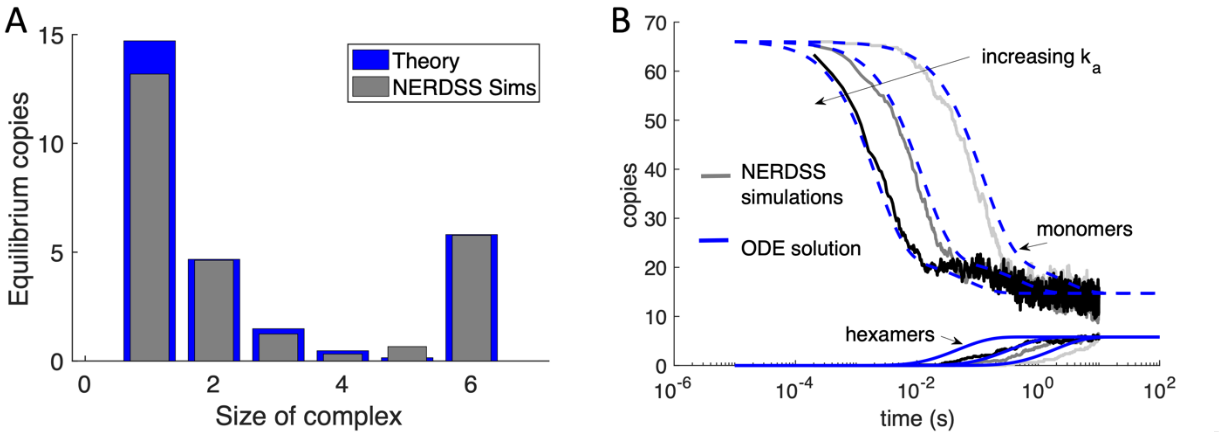


**Figure S2. NERDSS simulations of purely hexamer assembly in 2D are validated against theory.** A) Here, the Gag monomers could only bind through their hexamer sites, and the dimer sites were turned off (but could still exclude volume). With 66 initial Gag monomer copies initialized irreversibly on the spherical surface, the monomers could assemble into complexes from dimers up through to completed hexamers purely in 2D. The equilibrium yield here calculated numerically (blue bars) which agrees very well with the simulated equilibrium (gray bars). The free energy is $\Delta G_{hex}$=-8.93 $k_BT$. When the hexamer loop closes, the strength of the final two bonds is slightly less than $2\Delta G_{hex}$, and instead is $2\Delta G_{hex}$+2.3 $k_BT$. We introduce this small penalty to mimic that the hexamer structure is not ideal. B) We compare the kinetics of assembly from the NERDSS simulations (gray solid lines) with a set of non-spatial ODEs for hexamer formation solved numerically in MATLAB. The monomer population decreases from 66 copies (upper curves), while the hexamers assemble to their equilibrium value of ~6 (lower curves). The free energy for all simulations is the same as described in (A), and the microscopic rates increase from right to left as $k_a^{3D}$=0.025, 0.25, and 2.5 $nm^3/\mu s$, with the length-scale from 3D to 2D set at the same value used in the full system as $h$=10nm. The microscopic dissociation rates thus also increase accordingly from 2 to 200 $s^{-1}$. Diffusion D=0.2 $nm^2/\mu s$ for each membrane-bound monomer, and the binding radius for the hexamer interaction is $\sigma$=0.418 nm The agreement between the NERDSS simulations and the ODEs are relatively strong, although the simulated hexamers assemble more slowly. This is in large part due to the excluded volume of the monomers (via their dimer sites) that slows the collisions between the reactive hexamer sites.

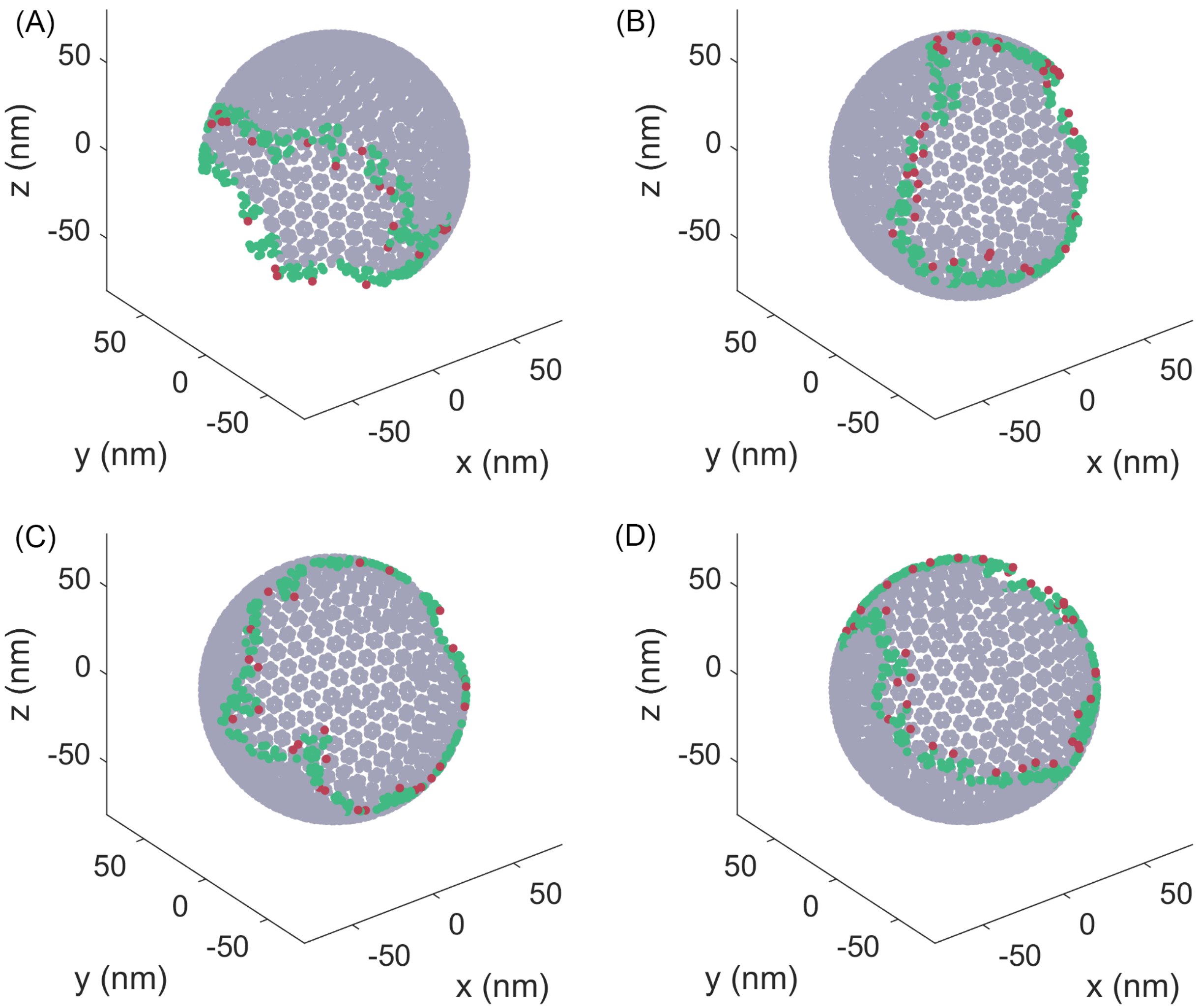


**Fig S3. Edge of the lattice.** Center-of-mass of each molecule is represented by a point. The green molecules are the molecules at the edge. A molecule is considered at the edge if it has less than 39 molecules within 15nm. The red molecules are the molecules that are single linked to the lattice.

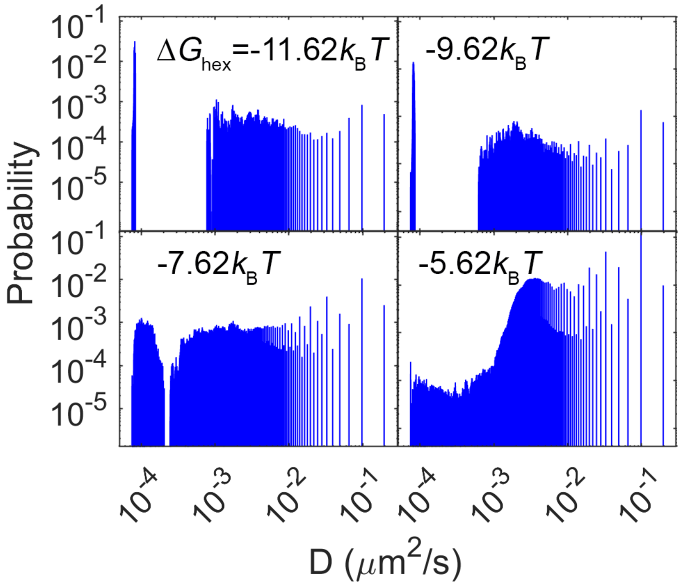


**Fig S4. Distribution of the diffusion constant of each molecule.** The most right bar is the molecule of monomers. The most left bar are the molecules within the largest complex. A clear separation of time-scales emerges as the lattice stabilizes due to the giant connected component. The rate constant for these simulations was the intermediate value of 0.025 $nm^2$/us.

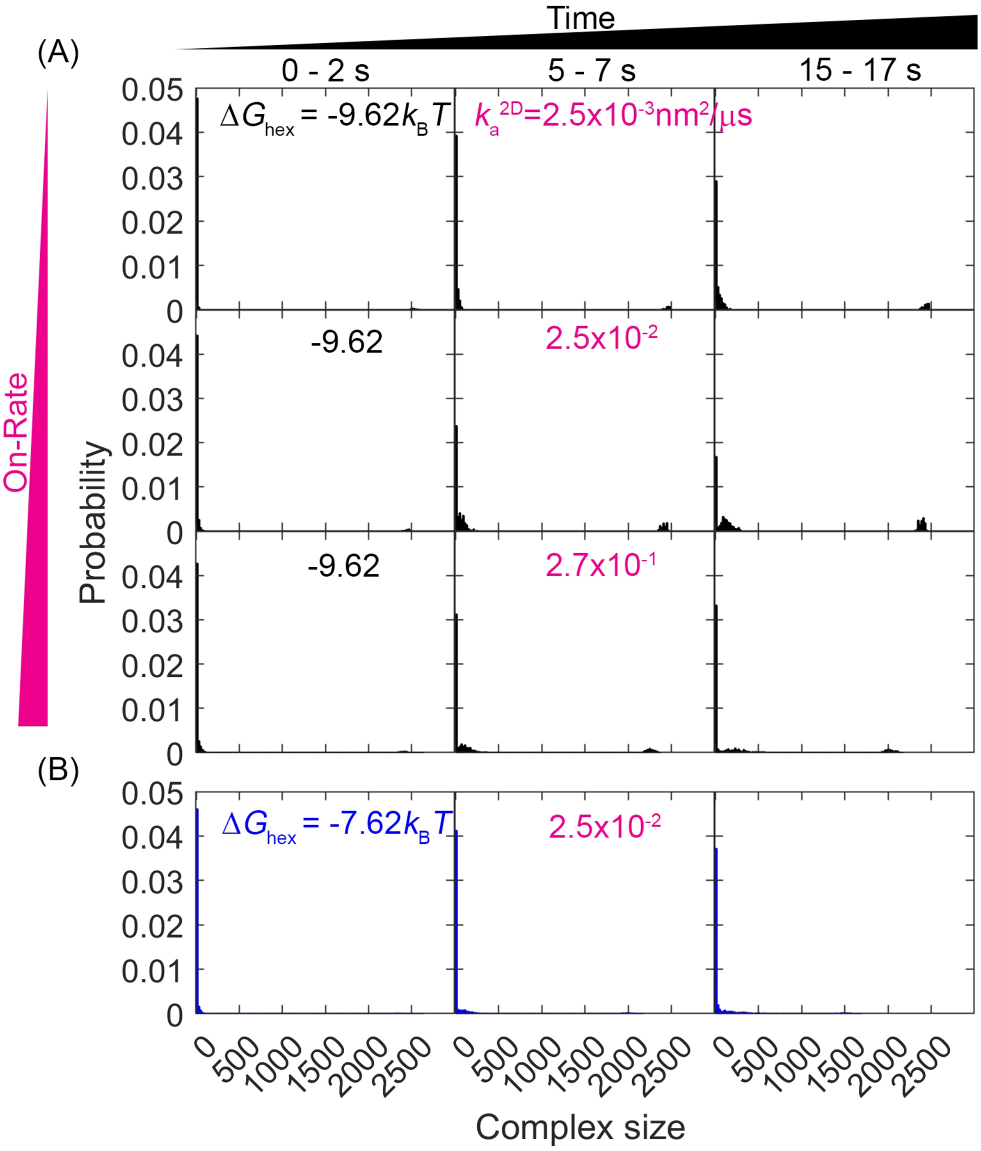


**Fig S5. Evolution of the lattice size distribution at different reaction rates and hexamer interaction strengths.** The data are the same as Fig3 in the main manuscript except that the plots extend the y-axis to 0.05.

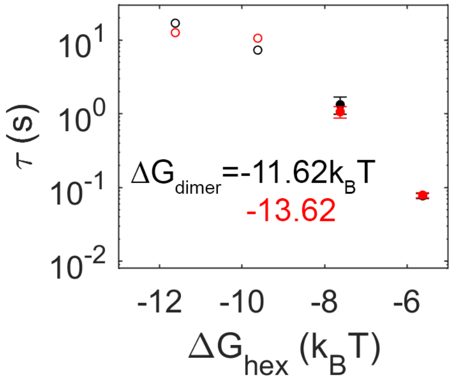


**Fig S6. Effect of dimer interaction strength of mean first-passage time.** Black data are from simulations with a dimer free energy of -11.62kBT, and red data are more stable at -13.62. Unlike the effect of changing the hexamer free energy, as evidence by the x-axis, a more stable dimer interaction has minimal effect on the MFPT. Filled data are from models where all trajectories produced dimerization events, open circles are lower bounds as not all trajectories produced events over the simulation time.

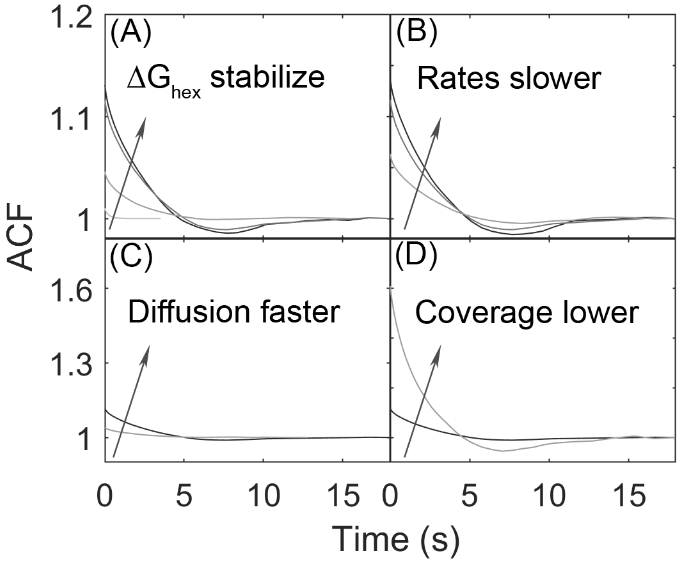


**Fig S7. Auto-correlation functions (ACF) at different free energies, reaction rates, diffusion, and surface coverage.** A) The ACF increases in the beginning as the hexamer energy stabilizes. B) The ACF increases in the beginning as the reaction rates become slower. C) The ACF increases as the diffusion becomes faster. D) The ACF increases as the surface coverage becomes lower.

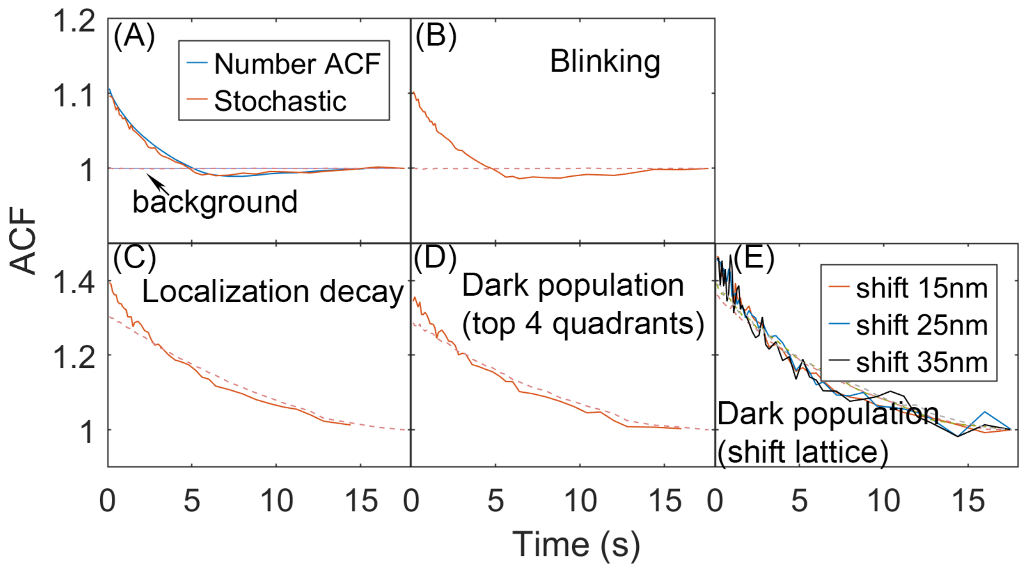


**Fig S8. Effects of introduced noise on the autocorrelation functions calculated from stochastic localization measurements on simulation trajectories.** A) From simulation we can directly count the number of monomers in each quadrant, and generate the complete number ACF. We can also perform a stochastic localization experiment, to mimic experiment, producing excellent agreement. In each frame, a monomer was localized here with 60% probability. Reducing the probability increases the noisiness of the ACF, but not its amplitude or timescales. No other 'error' was introduced into the localization measurement. For all plots, the background signal is shown in dashed red. The background is the correlation of the total copies counted across the full surface (no separation into quadrants). B) For the stochastic localization, we add blinking of the fluorophore. Each molecule, once localized, can be localized again within the 10 frames since its first localization, with maximal 3 total localizations. Each molecule is on average localized twice. C) The probability of localizing a molecule decays with time, such that early on, more localizations occur than later on in the trajectory. D) The surface of the sphere is assumed to be only partially 'visible' to the laser. Here only the top 4 quadrants are detected and analyzed, and the bottom 4 quadrants are 'dark'. Those monomers in the bottom half become visible once they diffuse into the top hemisphere. E) Similar to (D), except here the visible part of the sphere is asymmetric. The lattice is shifted to a distance along the x-axis and z-axis and only the molecules whose distance to the origin is less than $R_{sphere}$=67nm are visible. Monomers outside of that region are 'dark', until they diffuse into the visible part of the surface.
